# *Mycobacterium tuberculosis*-specific CD4 T cells expressing transcription factors associate with bacterial control in granulomas

**DOI:** 10.1101/2022.11.30.518638

**Authors:** Nicole L. Grant, Kristen Kelly, Pauline Maiello, Helena Abbott, Shelby O’Connor, Philana Ling Lin, Charles A. Scanga, JoAnne L. Flynn

**Author notes:** Corresponding Author: JoAnne L. Flynn, Address: University of Pittsburgh School of Medicine, 5058 Biomedical Science Tower 3, 3501 Fifth Avenue, Pittsburgh PA, 15261.

## Abstract

Despite the extensive research on CD4 T cells within the context of *Mycobacterium tuberculosis* (Mtb) infection, few studies have focused on identifying and investigating the profile of Mtb-specific T cells within lung granulomas. To facilitate identification of Mtb-specific CD4 T cells, we identified immunodominant epitopes for two Mtb proteins, Rv1196 and Rv0125, using a Mauritian cynomolgus macaque model of Mtb infection, providing data for the synthesis of MHC Class II tetramers. Using tetramers, we identified Mtb-specific cells within different immune compartments post-infection. We found that granulomas were enriched sites for Mtb-specific cells and that tetramer^+^ cells had increased frequencies of the activation marker CD69, and transcription factors T-bet and RORγT, compared to tetramer negative cells within the same sample. Our data revealed that while the frequency of Rv1196 tetramer^+^ cells was positively correlated with granuloma bacterial burden, the frequency of RORγT or T-bet within tetramer^+^ cells was inversely correlated with granuloma bacterial burden highlighting the importance of having activated, functional Mtb-specific cells for control of Mtb in lung granulomas.

## INTRODUCTION

*Mycobacterium tuberculosis* (Mtb), the causative agent of tuberculosis (TB), remains a global health problem despite over a century of observations and research. The bacterium itself is a complex organism with roughly 4,000 genes and an estimated 3,924 proteins, of which 204 are predicted to be secreted (1). Despite this large antigenic potential, immunodominant antigens overlap between human and macaque species (2). Several immunodominant antigens to Mtb have been studied with some being incorporated into vaccines currently in clinical trials, although the functions and relevance to virulence for many of these proteins remain incompletely understood. The immunodominant antigens included in several vaccine candidates consist of the well-known secreted proteins ESAT-6 and CFP-10, as well as the serine protease Rv0125, and the functionally less understood protein Rv1196 (3–9). CD4 T cells have been extensively studied in the context of Mtb infection, with critical functions related to their production of pro-inflammatory cytokines and interactions with other cells, including CD8 T cells and macrophages (10, 11). While most studies investigating CD4 T cells in TB disease use Mtb peptide stimulation, tetramer staining in blood, or single cell flow or RNA sequencing analysis, few provide insight into the presence and function of Mtb-specific CD4 T cells in tissue sites of infection such as lung granulomas, particularly in non-human primate (NHP) models (12–16).

While the large antigenic repertoire of Mtb could contribute to enhanced pathogen recognition by host T cells, it may also contribute to the difficulty in its clearance and complexity of immune responses as some antigens may act as “decoys” evading host protective responses (17). Following infection, granulomas are formed through the migration of innate and adaptive cells in response to the initiating bacillus, contributing to the overall microenvironment (18, 19). Previous studies investigating T cells in granulomas provide evidence that while they make up roughly 30-40% of all cells, only a fraction produce pro-inflammatory cytokines (20). Hypotheses as to why T cells may not produce high frequencies of pro-inflammatory cytokines within lung granulomas include, but are not limited to: 1) T cells in lung granulomas experience constant stimulation by antigen and thus become functionally exhausted; 2) the granuloma is comprised of spatial cellular compartments, limiting interactions between APCs and specific T cells; and 3) non-specific T cells are recruited to lung granulomas and thus the functionality is dependent on antigen-specificity. Previous studies from our lab and others indicate that there are low levels of multiple exhaustion markers on granuloma T cells and have predicted how the cellular spatial environment of the granuloma contributes to macrophage:T cell interactions (17, 21–23). However, the difficulty in studying antigen specific T cells in granulomas, which often have limited cells for analysis, presents a challenge to test these possibilities. Tools to identify Mtb-specific T cells within the granuloma would provide valuable insight into the phenotype and functionality of these cells.

Tetramers are a valuable tool for identifying and investigating functional attributes of antigen-specific cells, however, given the wide polymorphic diversity in MHC I and II molecules in humans and commonly used NHP models, designing useful tetramers can be challenging (24)(25–33). Mauritian cynomolgus macaques (MCMs), provide a useful model to develop and use tetramers to study antigen-specific cells as these NHPs emerged from a small, isolated, founder population, leading to a substantially reduced MHC diversity (29, 34, 35). The diversity in MHC I and II molecules in MCMs has been well studied, providing evidence of 7 distinct groups based on the expression of major MHC haplotypes (35). Using MCMs to include only those from the a specific haplotype (e.g. M1/M1) greatly reduces the major MHC I and II allele variation (35). MCMs have been developed as a model for Mtb infection, presenting with granulomas and other pathologies similar to humans and other NHP species, with a higher susceptibility to active TB disease similar to that of rhesus macaques (36–38).

Here, we mapped dominant epitopes for two immunodominant Mtb proteins in M1/M1 MCMs and acquired tetramers for these antigens. Using these new tetramers, plus two tetramers previously developed to identify CD4 T cells specific for CFP-10, we investigated Mtb-specific T cells within infected MCMs, providing insight into the presence of Mtb-specific T cells in the blood, BAL, lungs, lung granulomas, thoracic lymph nodes (LNs), and extra-pulmonary sites (EP) (3). Furthermore, we investigated the function of Mtb-specific CD4 T cells in lung lesions using transcription factors and activation markers and demonstrated an association between Mtb-specific CD4 T cells expressing T-bet or RORγT and a reduction in bacterial burden within granulomas.

## MATERIALS AND METHODS

### Ethics statement

All experiments, protocols, and care of animals were approved by the University of Pittsburgh School of Medicine Institutional Animal Care and Use Committee (IACUC). The Division of Laboratory Animal Resources and IACUC adheres to national guidelines established by the Animal Welfare Act (7 U.S. Code Sections 2131-2159) and the Guide for the Care and use of Laboratory Animals (Eighth Edition) as mandated by the U.S. Public Health Service Policy. Animals used in this study were housed in rooms with autonomously controlled temperature, humidity, and lighting. Most animals were pair-housed although some were singly housed based on temperament or uneven animal numbers in the study. Animals were provided with visual and tactile contact with neighboring conspecifics. Animals were provided water ad libitum, large biscuits specifically formulated for NHPs, and supplemented with pieces of fresh fruits and vegetable at least 4 days per week. An enhanced enrichment plan was implemented with three components: Encouragement of species-specific behavior involving toys and other manipulata that are filled with food treats, which are rotated on a regular basis. Puzzle feeders, foraging boards, and cardboard tubes encourage foraging behaviors. Adjustable mirrors accessible to the animals stimulate interaction between animals. Secondly, routine interaction between humans and macaques are encouraged. Interactions occur daily and consist of small food objects offered as enrichment and adhere to the established safety protocols. Animal caretakers are encouraged to interact with the animals while performing tasks in the housing area. Routine procedures are performed on a strict schedule to provide animals with a routine daily schedule. Third, all macaques are provided with a variety of visual and auditory stimulation including TV/video equipment playing cartoons for at least 3 hours per day which is rotated regularly so that enrichment is not repetitively played for the same group of animals as well as devices including food puzzles. All animals are checked twice daily and closely monitored to assess appetite, attitude, activity level, hydration status, etc. Following Mtb infection animals are monitored for evidence of disease (e.g. anorexia, weight loss, tachypnea, dyspnea, and coughing). Physical exams are performed on a regular basis. Ketamine, or other approved drugs, are used to sedate animals prior to all veterinary procedures (e.g. blood draws, bronchoalveolar lavage). PET-CT imaging was conducted every other week for this study and has proved very useful for monitoring disease progression. Veterinary technicians monitor animals closely for signs of pain or distress. If any signs are noted, appropriate supportive care is provided (e.g. dietary supplementation) and clinical treatments administered (analgesics). Any animal considered to have advanced disease or intractable pain or distress from any cause was sedated with ketamine and humanely euthanized using sodium pentobarbital.

### Animals, Mtb infection, and disease tracking by PET CT

Fifteen (10 for mapping and MHC allele testing and 5 for tetramer testing) Mauritian cynomolgus macaques (*Macaca fascicularis*) were obtained from Buckshire Corporation or Bioculture-Mauritius. Animals were placed in quarantine for either 30 or 60 days depending on the length of time in the United States and were monitored for signs of poor health and prior infection. After quarantine, the animals were transferred to the University of Pittsburgh Regional Biocontainment Laboratory BSL3 facilities and infected with a low dose of (<20 CFU) Mtb Erdman via bronchoscopic instillation as previously described (39, 40). Infection trajectory and disease was tracked using ^18^F-fluorodeoxyglucose (FDG) PET-CT for granuloma formation and lung inflammation every 4 weeks. PET CT scans were analyzed using OsiriX viewer with a 1mm limit of detection as previously detailed (41).

### Bronchoalveolar lavage (BAL)

Bronchoalveolar lavage was performed as previously described (40). In brief, a 2.5mm diameter bronchoscope was inserted into the trachea of a sedated animal and placed in the right middle of lower lobe, where a saline solution (40mL) was introduced and suctioned into a sterile 50mL conical tube. An aliquot was plated on 7H11 agar plates to determine CFU, which were counted after 3 weeks of incubation at 37°C/5%CO_2_. The remaining BAL fluid was centrifuged at 1800 rpm for 8 minutes at 4°C. Cells were resuspended in 1mL of sterile PBS and counted using a hemocytometer and aliquoted for use in tetramer and flow cytometry staining.

### Necropsy procedures

Necropsies were performed on animals at the pre-determined study endpoint or humane endpoint determined by clinical evaluation or PET CT scan. In brief, animals were sedated with ketamine, maximally bled and then euthanized with pentobarbital, and tissue samples extracted. PET CT guided necropsy procedures were followed as previously described in which each granuloma or disease pathology is identified on the scan and matched to the lung location for dissection (41). All lung tissues, thoracic lymph nodes, spleen and liver were also obtained. Individual samples were placed in RPMI 1640 media (Fisher, BW12-167F) and homogenized for single cell suspensions. Sections of lymph node and lung tissue samples were formalin fixed for histopathological analysis. Single cell suspensions were counted using a hemocytometer to determine viable cell counts and plated on 7H11 agar plates and incubated for 21 days at 37°C for CFU determination. To increase the likelihood of capturing tetramer^+^ populations, smaller samples were combined from the same animal to increase the number of cells stained for more distinct gating. Of note, granulomas from an individual animal were not combined with those from a different animal. Using this method, we were able to identify tetramer^+^ populations with >50 events in most of our pooled or individual granuloma samples, thus allowing sufficient cell numbers to analyze the functionality of tetramer^+^ CD4 T cells.

### IFN-γ ELISPOTS

Peripheral blood mononuclear cells (PBMCs) were isolated from blood draws pre- and post-Mtb infection. White membrane plates (Fisher Scientific, Cat. #MSIPS4W10) were prepared using 30% ETOH followed by three washes with sterile 1XPBS. Primary antibody (anti-IFN-γ clone MT126L) was used as a capture antibody and following addition, plates were incubated overnight at 37°C. Following addition of the capture antibody, ELISPOT plates were washed three times with sterile 1XPBS and blocked for two hours at 37°C using RPMI 1640 (Sigma, Cat. #R0883) supplemented with 1% L-glutamine (Thermo Fisher, Cat. #25030149), 1% HEPES (Thermo Fisher, Cat. #SH3023701), and 10% human A/B serum (Gemini Bio-Products, Cat. #100-512). Peptides were added to prepared plates at a final concentration of 1μg/mL-2μg/mL followed by 1x10^5^-2x10^5^ fresh or frozen (rested overnight) PBMCs. Plates were incubated for 48 hours at 37°C and immediately washed 6 times with 1XPBS. Following washing, a filtered secondary antibody (anti-IFN-γ, clone 7-B6) was added to plates and incubated for two hours at 37°C. Plates were washed 6 times with 1XPBS then followed by addition of a Streptavidin-HRP antibody for 45 minutes at 37°C. Lastly, an AEC substrate (Vector Laboratories, Cat. #SK-4200) was added according to manufacturer’s instructions for 5-8 minutes to allow plate to develop followed by 3 washes each of diH20 and 1XPBS. All samples were run in duplicate wells for each experiment and represented as the average spot forming unit (SFU). Plates were dried in a cool, dark location and read using an Immunospot ELISPOT plate reader (Cellular Technology Limited, Cleveland OH, USA) and manually assessed for quality control.

### Peptide resuspension and mapping strategies

Sequences for Mtb proteins Rv1196 and Rv0125 were obtained from publicly available databases and peptide libraries were generated using a commercially available peptide library design tool (PEPscreen, Millipore Sigma, Merck, Darmstadt, Germany) with peptide lengths of 20 amino acids (a.a.) and an overlap of 10 a.a. Individual peptides were ordered from Genscript and resuspended in DMSO (with a calculated final concentration of no greater than 10%) and sterile 1X PBS according to molecular weight to obtain a stock concentration of 10mM. Stock concentrations were diluted using sterile 1X PBS to 1mg/mL working concentrations and peptides were used at 1-4μg/mL according to assay standards.

### Plasmid generation and purification

Plasmids containing major M1 allele variants (both MHC I and MHCII) (Supplementary Figure 1A) were kindly gifted from collaborators at the University of Wisconsin (O’Connor lab). To obtain larger stocks, plasmids were transformed into E. coli using heat shock transformation, selected on antibiotic plates and grown in liquid media. Plasmids were purified from the E. coli strains with a Qiagen miniprep kit using a vacuum manifold. Concentrations of plasmids were determined using a nanodrop reader with OD 260/280 ratios at ∼1.8 considered pure.

### Transfection of mammalian cells for allele expression

Several transfection methods were tested and success rates varied based on the plasmids containing allele variants. Mammalian cells lacking MHC I (K562 cells) or MHC II (RM3 cells) proteins (kindly sent by Shelby O’Connor) were used for allele restriction determination. Electroporation using a BTX ECM630 was determined as the optimal system for transfecting RM3 cells containing MHC II allele variants under the following conditions: mode: low voltage, capacitance 250μF, resistance: none, charging voltage: 200 V, Chamber BTX disposable cuvette (2mm gap), field strength: 750V/cm, sample volume: ∼300μL (3, 42–44). Prior to electroporation RM3 cells were kept in a log growth phase by subculturing and observing flasks for cell health and debris. Plasmids were added to cuvettes (10μg of total plasmid; e.g. 5μg of DRA and 5μg of DRB) along with RM3 cells and resuspended in 150mL of ECM buffer. Following electroporation, RM3 cells were grown under unsupplemented media for 24 hours at 37°C, 5%CO_2_. After 24 hours, media was changed and cells were grown in R10 media (RPMI 1640 + 10% FBS, 1% antimycotic/ antibiotic) for 48 hours. At this time, hygromycin B (Sigma, Cat. #H0654-500mg) was added to the media at a final concentration of 400μg/mL for an additional six days, with sub-culturing every 48-72 hours. Transfectants were stained for viability using a Live dead blue dye followed by anti-HLA DR DP DQ antibodies (BioRad clone Bu26, Cat. #MCA2497F; Beckman clone I3, Cat. #CO6604366; BioLegend, clone Tü39, Cat. #361704) or anti-HLA DP (Leinco, clone B7/21, Cat. #H129) for detection of MHC II proteins.

### Generation of EBV transformed-B lymphoblastic cell lines

Autologous BLCLs were generated using isolated PBMCs from pre-infection blood draws for use as antigen presenting cells and culturing and testing T cell lines. Isolated PBMCs (1x10^5^-2x10^5^) were added to 96 well round bottom plates with 50μL of *Herpesvirus papio* supernatant, from filtered supernatants of S594 cells, and grown in R20 (RPMI + 20% FBS). Media was changed 3-4 days following addition of S594 supernatant and thereafter every 2-3 days. When cells started to clump and media was slightly yellow, the BLCLs were split into additional wells of the 96 well plate. Cells from 96 wells were combined into wells of a 24 well plate in R20 and expanded into larger size wells (6-well plate) with T25 or T75 flasks being seeded for freezing or for use in experiments.

### Peptide specific T cell generation and expansion

A series of experiments were performed to generate T cells for long term culture and testing according to protocols provided by the O’Connor lab and using published protocols (45). However, in our experiments, stimulation with peptide-pulsed irradiated BLCLs did not yield stable peptide-responsive T cell lines. We used the following approach for the data provided here: Isolated PBMCs were plated in 12 well or 6 well plates at a concentration of 1x10^6^ cells/mL in RPMI media supplemented with 15% FBS, IL-2 (Abcam, Cat. #ab119439, 1μL/10mL) and IL-7 (Abcam, Cat. #ab73201, 1μL/20mL). Peptide of interest was added to individual wells at a final concentration of 1μg/mL peptide. Cells were incubated at 37°C, 5%CO_2_ for 48 hours and then washed with sterile 1XPBS and rested for an additional 4 days in R15+IL2/IL7 media.

### Peptide specific T cell allele restriction testing

PBMC T cell cultures expanded with the dominant epitope for RV1196 or Rv0125 were tested for specific allele presentation using RM3 cells transfected with MHC II alleles at a 1:1 or 2:1 ratio (i.e. T cell:transfected RM3 cell)(Supplementary Figure 1). Transfected RM3 cells were prepared for T cell testing by adding 1μg/mL peptide to individual wells of a 96 well plate and incubated for 90 minutes at 37°C, 5%CO_2_. Following incubation, prepared T cells were added to each well of a 96 well plate and incubated for another 12 hours in the presence of a Golgi plug inhibitor, BFA (BD Biosciences, Cat. #555029). Stimulation plates were spun down and washed with 1XPBS following 12-hour incubation. Flow cytometry staining was performed for viability (Live/dead fixable blue dead cell stain kit, Thermo Fisher, Cat. #L34962) and surface marker expression for CD3 (BD, clone SP34-2), CD4 (BD, clone L200), CD8 (BD, clone RPA-T8), HLA DR/DP/DQ (BioRad clone Bu26; Beckman clone I3; BioLegend, clone Tü39), and intracellular cytokines IFN-γ (BD, clone B27), and TNF (BD, clone Mab11) using standard flow cytometry and intracellular staining protocols. Samples were run on a BD LSRII or a Cytek Aurora (Cytek, Bethesda, MD, USA) and analyzed using FlowJo software (BD, version 10).

### Tetramerization

Monomers were prepared by the NIH tetramer core as requested for peptide and allele sequences. Once received, samples were aliquoted and stored at -80°C until time of use. To tetramerize monomer, fluorescently labeled streptavidin (SAV) was added in 10-minute increments, 10 times based on the concentration of the SAV as previously described in the NIH tetramer core published protocols (46). Serial addition of SAV allows for binding of free monomer while leaving little excess SAV. After tetramerization, staining was performed on frozen PBMCs or expanded T cells to ensure proper complex formation. Tetramers were stored at 4°C and used prior to 2 months from synthesis to allow for minimal tetramer disaggregation. Tests were performed to determine the optimal amount of tetramer to use per sample, by comparing control tetramer staining to Rv1196_371-385_ tetramer staining, showing that 4ug/mL was adequate to visualize a distinct population of tetramer^+^ cells. For tetramer staining, samples were prepared and stained in a 500nM RPMI+dasatinib (Fisher, Cat. #NC0897653) solution and washed with 50nM dasatinib solution in 1XPBS. Samples were stained with tetramer, control tetramer (CLIP), or no tetramer for 30 minutes at room temperature (RT) followed by two washes with 50nM 1XPBS. To maximize the use of validated fluorophores for tetramer conjugation, we conjugated monomers for CFP-10_36-48_ and CFP-10_72-85_ in the same fluor (BV421), since both detect T cell responses to the protein CFP-10, and conjugated monomer for Rv1196_371-385_ with PE and the monomer for Rv0125_81-92_ in APC. The concentration of tetramer needed for clear identification of tetramer^+^ populations was determined using expanded T cell lines and compared to control tetramer staining at the same concentration. Granuloma lung samples were stained with control tetramers (CLIP monomer provided by NIH and tetramerized by us), no tetramer, no transcription factor, or tetramer plus transcription factors, depending on cell numbers, with all the controls being performed for each animal to ensure accurate gating for analysis of tetramer^+^ and transcription factor^+^ cells.

### Flow cytometry and transcription factor staining

Bronchoalveolar lavage was performed as described above and cells counted for staining at 3- and 4-weeks post infection for all animals and 8 weeks for 1 animal. Similarly, necropsy tissues were homogenized, counted, and single cell suspensions prepared by washing with 500nM dasatinib RPMI solution. After tetramer staining and washes were performed, cells were stained with Zombie near infrared (NIR) (BioLegend, Cat. #423105) diluted 1:1000 per manufacturer’s recommendation for 10 minutes at 4°C in the dark. Cells were washed with 50nM dasatinib in 1XPBS twice and spun down each time at 2000 rpm, for 3 minutes at 4°C. Following washes, cells were stained with surface antibody cocktail diluted in 50nM dasatinib FACS (1XPBS + 5% FBS) buffer for 20 minutes at 4°C in the dark. Cells were washed twice using 50nM dasatinib FACS solution, spun as described above, and fixed for a minimum of 10 minutes in 4% paraformaldehyde (PFA) for processing outside of the BSL3. Samples were washed in 1XPBS and placed at 4°C overnight for continued staining the following day. To begin staining the following day, samples were spun at 2000 rpm for 3 minutes at 4°C and supernatants decanted. True nuclear transcription factor buffer kit (BioLegend, Cat.#424401) was used for detection of transcription factors within cells as per the manufacturer’s recommendation. Following a 1 hour permeabilization at RT in the dark and two subsequent washing steps, samples were stained for 1 hour at RT in the dark using antibodies targeting transcription factors, granzyme B, and CD69. Samples were washed twice using 50nM dasatinib FACS solution and resuspended in 1XPBS. Granuloma lung samples were stained with control tetramers (CLIP monomer provided by NIH and tetramerized by us), no tetramer, no transcription factor, or tetramer plus transcription factors, depending on cell numbers, with all the controls being performed for each animal to ensure accurate gating for analysis of tetramer^+^ and transcription factor^+^ cells. For analysis, we employed a gating strategy using an inclusive CD4 gate (gated on all CD4^+^ cells, including CD4^+^CD8^+^) to optimize our ability to identify the tetramer^+^ cells (Supplementary Figure 2). Transcription factor analysis was performed on tissue samples where there were at least 50 events in the parent (i.e. Tetramer+) gate. A Cytek Aurora was used to acquire sample events and analysis was performed on unmixed data files (adjusted for reference controls) using FlowJo version 10.

### Statistical Analysis

Statistical analysis was performed in GraphPad Prism (version 9). Data were tested for normality using the Shapiro-Wilk test. Groups were compared using a Wilcoxon ranked sum analysis on paired samples or a Mann-Whitney on unpaired samples. For correlations, CFU (+1) was log_10_- transformed and tested for normality. For normal data, Pearson r correlation coefficient was reported; otherwise, Spearman r correlation coefficient was reported. P values < 0.05 were considered significant.

## RESULTS

### Identifying a dominant epitope for Rv1196 and Rv0125 in M1/M1 MCMs

A total of 10 Mtb-infected MCMs (including M1 homozygous and M1 heterozygous animals) were used to epitope map two Mtb proteins, Rv1196 and Rv0125, using IFN-γ ELISPOTs (Figure 1). To investigate the dominant epitope binding region for these proteins, we used PBMCs from various timepoints throughout infection, with PBMCs from 4-8 weeks post-infection (p.i.) eliciting the best overall IFN-γ responses. To narrow down the dominant epitope binding region individual 20 amino acid (a.a.) peptides with overlapping 10a.a. regions were pooled in a matrix format (Figure 1A, D, Supplementary Tables 1 and 2).

**Figure 1:**
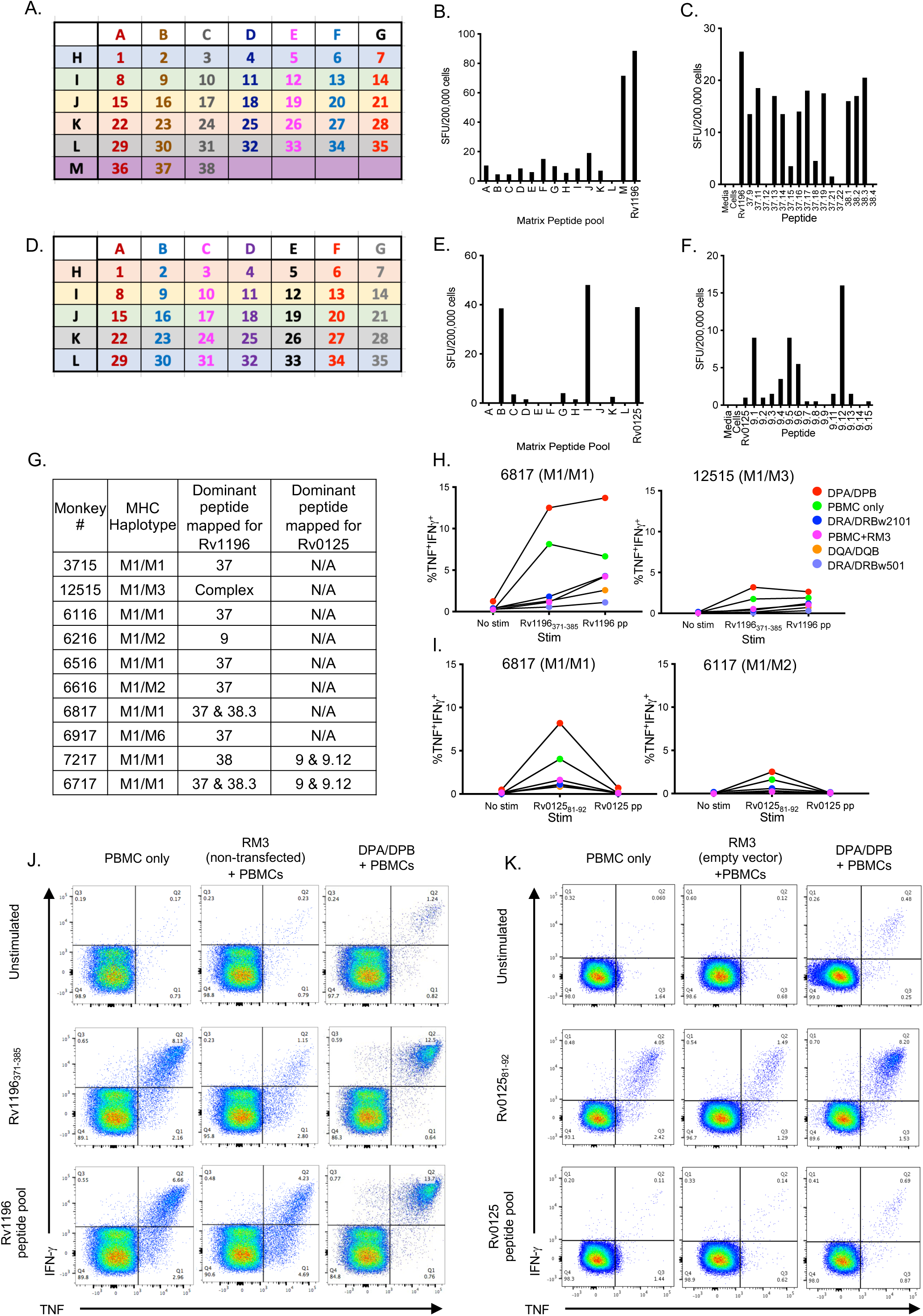
Epitope mapping and MHC allele restriction for Mtb proteins Rv1196 and Rv0125. (A) Peptide pools following a matrix mapping strategy were generated for Rv1196 where letters indicate peptide pools and numbers indicate individual 20 amino acid peptides (Supplementary Table 1). IFN-γ ELISPOT using stimulated PBMCs with Rv1196 peptide pools from a matrix pooling strategy (B) and truncated and adjacent peptides (C). (D) A similar approach was performed for Rv0125 peptide pools following a matrix mapping strategy for Rv0125 (Supplementary Table 2). IFN-γ ELISPOT using stimulated PBMCs peptide pools from a matrix pooling strategy (E) and truncated and adjacent peptides (F). (G) List of Mtb-infected MCMs used for epitope mapping for Rv1196 and Rv0125. (H, I) T cells were expanded from PBMCs isolated from MCMs and co-cultured for 12 hours with Rv1196_371-385_ peptide (I) or Rv0125_81-92_ peptide (J) and stimulated with RM3 cells transfected with different M1/M1 MHC alleles (color legend shown) and flow cytometry performed to determine the MHC II presenting allele. The highest frequency of TNF^+^IFN-γ^+^ production from CD4^+^ cells was observed when cultured with either peptide and RM3 cells transfected with the DPA/DPB allele. (J, K) Representative flow cytometry plots showing expanded T cells co-cultured with DPA/DPB-transfected (or untransfected) RM3 cells and unstimulated or stimulated with Rv1196_371-385_ or Rv1196 peptide pool (J) or Rv0125_81-92_ or Rv0125 peptide pool (K) for animal #6817 (M1/M1).

For mapping Rv1196, the highest IFN-γ responses were to peptides #37 and 38, indicating that the optimal binding region likely included the overlapping 10 a.a. region (Figure 1B). To further evaluate the amino acids involved in binding, we used truncated and shifted peptides of 9-20 a.a. lengths in PBMC IFN-γ ELISPOTs (Figure 1C and Supplementary Table 1). For Rv1196, we identified the optimal binding region to be between amino acid 371-385 (peptide Rv1196 38.3)(Figure 1G). Only peptides containing the VM (valine-methionine, a.a. 383 and 384) elicited IFN-γ responses, suggesting that VM is critical in peptide binding to M1 MHC II molecules (Supplementary Table 1).

A similar approach was taken using peptides for Rv0125 with the highest response elicited from Rv0125 peptide 9 (Rv0125_81-100_) and by truncated peptide 9.12 (Rv0125_81-92_) (Figure 1D-G and Supplementary Table 2). Results from IFN-γ ELISPOTs revealed that the dominant epitope binding region for Rv1196 and Rv0125 were 15 and 12 a.a., respectively. As MHC I binds peptides of smaller sizes, it was likely that these peptides are bound by MHC II and recognized by CD4 T cells, though based on size, Rv0125_81-92_ is small enough to be bound by MHC I (47).

### Rv1196_371-385_ and Rv0125_81-92_ are presented by the DPA/DPB allele

There are two main steps in the tetramer design process: (1) identifying the dominant epitope as outlined above and (2) identifying the MHC and allele presenting the dominant peptide to either CD8 or CD4 T cells (MHC I or MHC II, respectively). This critical step involves generating T cell lines, APC lines, and cell lines expressing specific MHC alleles. The generation of T cell lines has been described in humans, mice and NHP using media supplemented with recombinant IL-2 and IL-7 (48, 49). Although it has been reported that T cell lines can be passaged 10 weeks or more and still be functional, we did not observe cytokine production in our T cell lines grown and stimulated for more than 3 weeks in culture with irradiated BLCLs and peptide (50). We developed a protocol for expanding T cells based on protocols provided by University of Wisconsin. Following a 12-hour co-culture experiment, in which our expanded T cell lines were incubated with autologous peptide stimulated PBMCs, we performed flow cytometry staining for surface markers (CD3, CD4, and CD8) and cytokines (IFN-γ and TNF) to determine whether the peptides were presented to CD4 or CD8 T cells. This demonstrated that CD4 T cells recognized and produced IFN-γ and TNF in response to both peptides (Rv1196_371-385_ and Rv0125_81-92_) and therefore were likely MHC II restricted (Supplementary Figure 1B).

These data focused our efforts on producing M1-specific MHC II allele expressing RM3 cells, a cell line lacking MHC II expression (see Materials and Methods). We generated four sets of RM3-transfected cells representing the dominant M1/M1 DR, DP, and DQ alleles (Supplementary Figure 1A). There was variable frequency in allele expression despite within protocol consistency, with DP and DR alleles exhibiting lower frequencies of expression (Supplementary Figure 1C). After testing an alternative set of antibodies (different clones) for the detection of MHC II proteins we observed a higher frequency of allele expression, indicating that our transfection experiments were successful and that available anti-DR/DP/DQ antibodies do not detect all alleles equally (Supplementary Figure 1D).

RM3-allele expressing cells (RM3-DPA/DPB, RM3-DQA/DQB, etc.) were co-cultured with expanded peptide-stimulated T cells to identify the specific M1 MHC II allele presenting Rv1196_371-385_ or Rv0125_81-92_. The frequency of CD4 T cells producing IFN-γ and TNF was evaluated using flow cytometry staining in each of the following conditions: (1) PBMCs alone; (2) RM3 cells with empty vector; (3) RM3-DRA/DRB*w501; (4) RM3-DRA/DRB*w2101; (5) RM3-DQA/DQB; or (6) RM3-DPA/DPB in the presence of Rv1196_371-385_ or Rv0125_81-92_. This experiment was performed using frozen or fresh PBMCs from two Mtb-infected M1 MCMs for each specific peptide and revealed that both Rv1196_371-385_ and Rv0125_81-92_ are presented by the major DPA/DPB allele (Fig 1H-K). This information was provided to the NIH tetramer core where tetramers and monomers were prepared for DPA/DPB Rv1196_371-385_ and DPA/DPB Rv0125_81-92_ and two previously published CFP-10 tetramers (DRA/DRB w501 CFP-10_36-48_ and DRA/DRBw501 CFP-10_71-85_) (3) for interrogating samples from Mtb infected MCMs.

### Mtb-specific CD4 T cells can be observed in airways and blood of infected macaques

To gain insight into the presence and function of Mtb-specific cells in M1 homozygous MCMs, 5 animals were infected with a low dose of Mtb and monitored throughout infection using PET CT imaging as previously described (41). Previous studies showed that MCMs are more susceptible to Mtb infection than Chinese cynomolgus macaques, therefore the study endpoint was planned for 10 weeks post-infection (36). However, Mtb disease progressed faster in 3 of the animals, resulting in a clinical endpoint and necropsy at 6-9 weeks post-infection (Supplementary Figure 2A).

Mtb-specific CD4 T cells in the blood were tracked throughout infection by monitoring IFN-γ responses following PBMC stimulation with dominant peptides (Figure 2A-C); all animals responded to CFP10, Rv0125, and Rv1196 dominant peptides, although responses were variable across animals and time points. At necropsy, PBMCs were stained with an optimized flow cytometry panel including 4 tetramers specific for the 3 Mtb antigenic targets (CFP-10, Rv1196, and Rv0125) (Figure 2D, Supplementary Figure 2D). Although tetramer^+^ cells represent a low frequency of CD4 T cells in blood of infected macaques (0.1-0.18%, medians), a distinct population of Mtb-specific cells was observed in the blood at necropsy (Supplementary Figure 2D).

**Figure 2:**
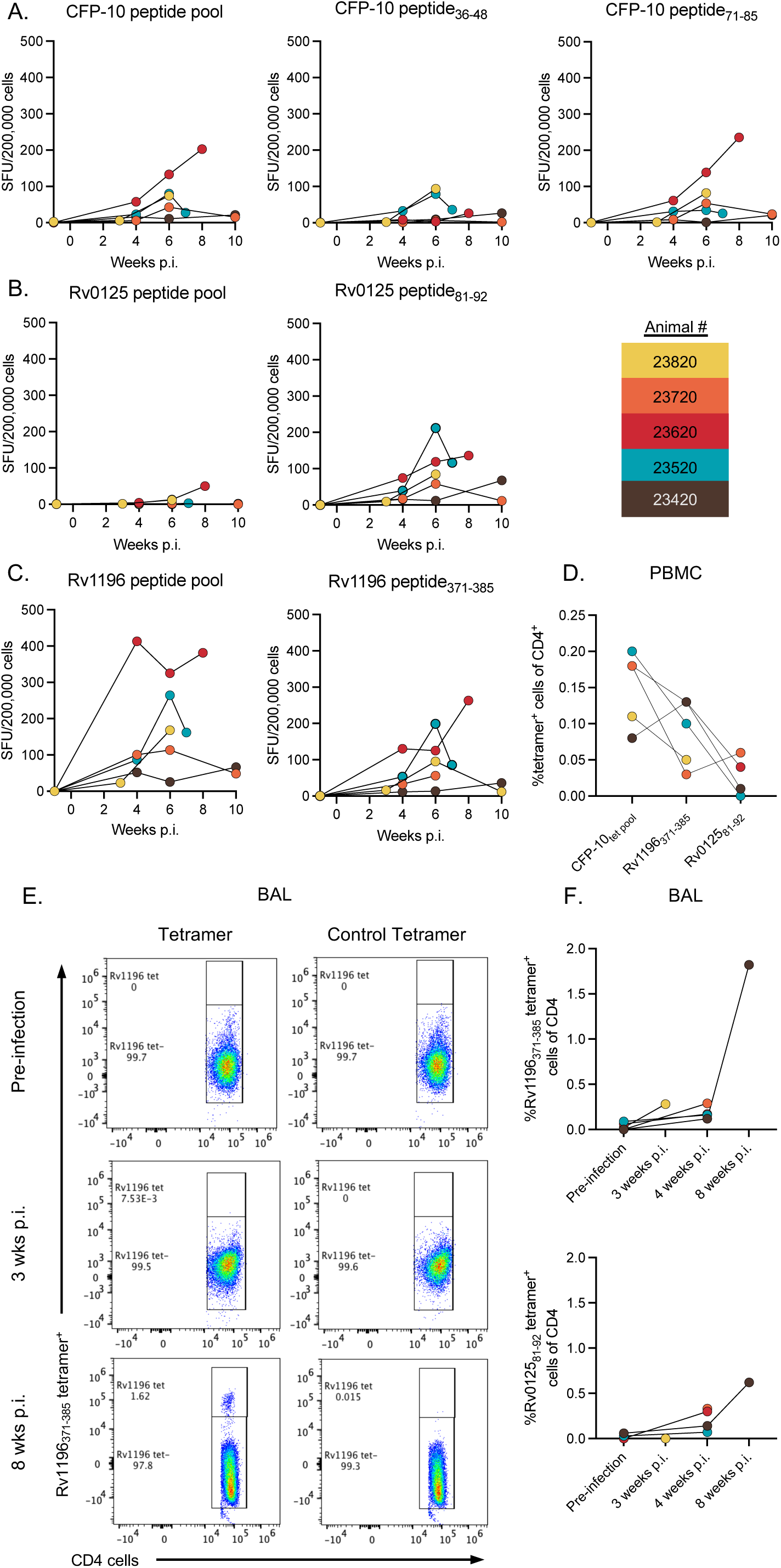
Identifying tetramer^+^ cells in the blood and BAL. IFN-γ ELISPOT response to (A) CFP-10 peptide pool, CFP-10_36-48_, and CFP-10_71-85_, (B) Rv0125 peptide pool and Rv0125_81-92_, and (C) Rv1196 peptide pool, and Rv1196_371-385_ in PBMCs throughout Mtb infection in 5 MCMs (weeks post-infection; color legend shown). (D) Frequency of tetramer^+^ cells of CD4^+^ cells in PBMCs at necropsy. (E) Flow cytometry plots showing control tetramer or Rv1196_371-385_ tetramer staining, gated on CD4 T cells, in pre-infection, 3, and 8 weeks post infection BAL from animal #23420. (F) Frequency of Rv1196_371-385_ tetramer^+^ CD4+ T cells (top) and Rv012581-92 (bottom) in BAL before and during infection.

To monitor the appearance of tetramer^+^ cells in the airways, flow cytometry was performed on cells obtained by BAL beginning at 3-4 weeks and in one animal at 8 weeks post infection. There was an increase in the frequency of tetramer^+^ cells as infection progressed (Figure 2E, F). The adaptive immune response to Mtb is slow to evolve in macaques (and humans) which could account for the low frequency of Mtb-specific T cells in airways at 3 and 4 weeks (51). Given the low numbers of tetramer^+^ cells in BAL samples, further analysis on cell function was not performed.

### Identifying Mtb-specific CD4 T cells in thoracic lymph nodes, lungs and granulomas

One of the gaps in knowledge is the frequency and functionality of Mtb specific T cells within lung, lymph nodes (LNs), and granulomas in NHPs or humans. The primary site of Mtb infection is the lung granuloma, however, a significant amount of disease including the formation of granulomas occurs within thoracic LNs (52). In this study, the median bacterial burden for involved LNs, i.e. those that were CFU positive, was higher than the median CFU for lung lesions (granulomas, clusters, and consolidations) (median for involved LN: 1.65x10^4^ CFU, median for lung lesions: 6.5x10^2^) (Figure 3A). Given the high bacterial burden and presence of granulomas in involved LNs, we hypothesized that Mtb tetramer^+^ cells could also be found within these sites. Using thoracic LN samples isolated at necropsy, we were able to identify Rv1196 tetramer^+^ cells within both involved LNs (CFU positive [CFU^+^] or gross detection of granuloma formation) and uninvolved LNs (CFU negative [CFU^-^] and lacking gross detection of granulomas) (Figure 3B). There was a range in frequency of Rv1196_371-385_ tetramer^+^ cells within thoracic lymph nodes between 0.0% and 0.21% with CFU^+^ LNs having a significantly higher frequency of Rv1196 tetramer^+^ cells compared to CFU^-^ LNs (CFU^+^ median: 0.073%, CFU^-^ median: 0.019%, p=0.0041) (Figure 3C, D). To identify the frequency of all tetramer^+^ cells in thoracic LN samples (i.e. using all 4 tetramers), we applied a Boolean OR gating strategy which combines the events from the individual tetramer^+^ gates; this was only performed on a subset of samples for which we were able to use all 4 tetramers. The frequency of all tetramer^+^ cells reflected the frequency ranges of Rv1196_371-385_ tetramer^+^ cells, providing a wider range for animal 23720, revealing that this animal either did not have many Rv1196_371-385_ tetramer^+^ cells or had poor Rv1196_371-385_ tetramer staining (Figure 3E). As with Rv1196_371-385_ tetramer^+^ cells, there were significantly higher frequencies of all tetramer^+^ cells in thoracic CFU^+^ LNs compared to thoracic CFU^-^ LNs (Figure 3F).

**Figure 3:**
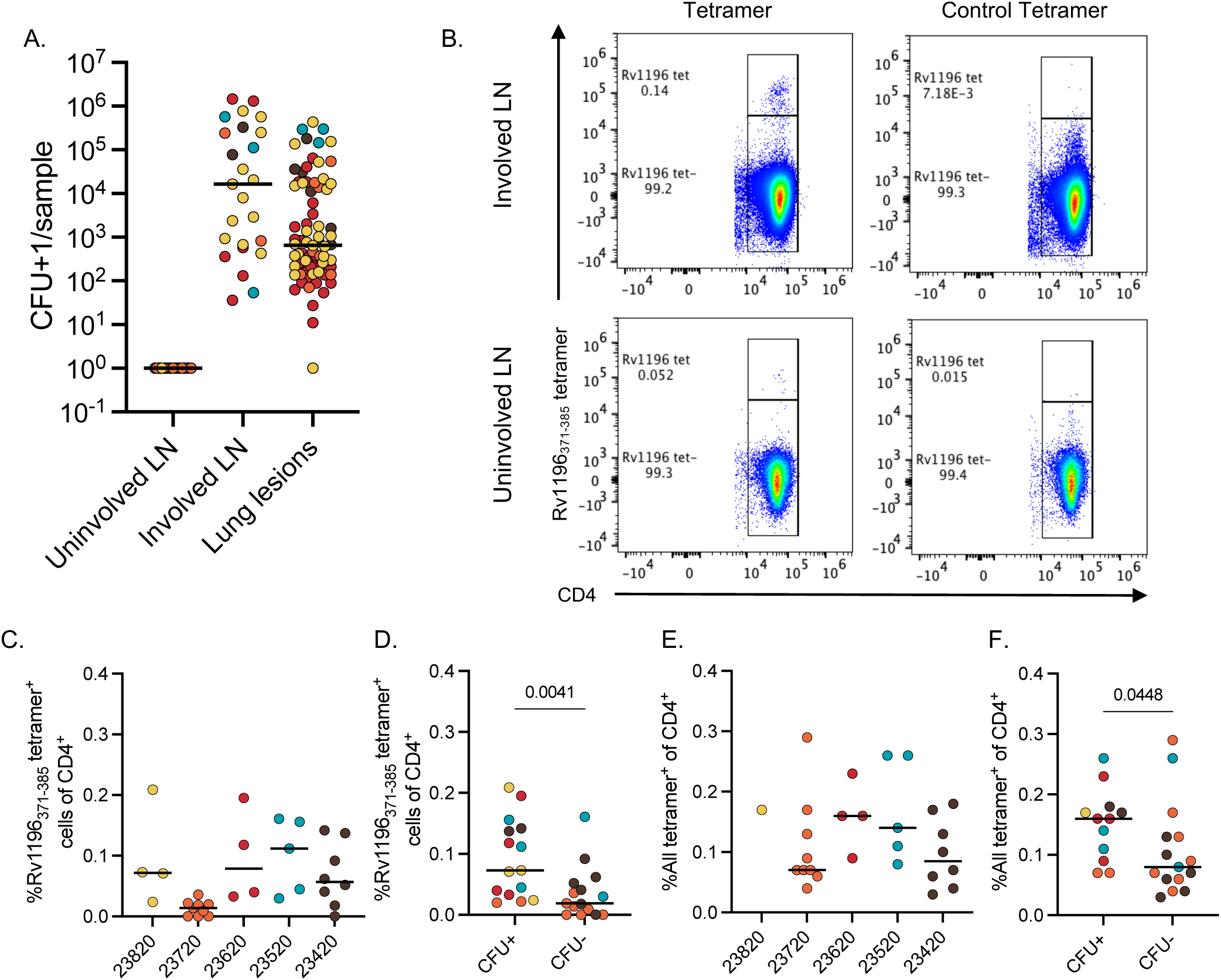
Detection of tetramer^+^ cells in thoracic LNs. (A) Mtb bacterial burden in uninvolved thoracic LNs (CFU negative or no gross granuloma detected), involved thoracic LNs (CFU positive or gross detection of granuloma), and lung lesions (individual granulomas, clusters, and consolidations), colored by animal (color legend as in Figure 2). (B) Example flow cytometry plots, identifying control tetramer^+^ or Rv1196 tetramer^+^ cells in involved thoracic LN and uninvolved thoracic LN (monkey 23420). (C) Frequency of Rv1196 tetramer^+^ cells of CD4 T cells in thoracic LNs by animal. (D) Comparison of Rv1196 tetramer^+^ cells of CD4+ T cells in CFU^+^ and CFU^-^ thoracic LNs. (E) Results of Boolean gating for all tetramer^+^ cells of CD4+ T cells in thoracic LNs by animal. Only samples for which all 4 tetramers were used are shown. (F) Comparison of all tetramer^+^ cells of CD4+ T cells in CFU^+^ and CFU^-^ thoracic LNs. Mann-Whitney tests performed to compare the medians of CFU^+^ vs CFU^-^ thoracic LNs (D and F). Points represent individual samples, colored according to animal where bars represent medians.

A total of 51 lesions (granulomas or granuloma clusters, individual or pooled) were analyzed for this study with a range in bacterial burden between 7x10^1^-4.3x10^5^ CFU (Figure 4A). The frequency of tetramer^+^ cells was variable in granulomas and within and across animals, with a median frequency of 0.83% of CD4 T cells staining positive for CFP-10 tetramers and median frequency of 1.27% of CD4 T cells staining positive for the Rv1196_371-385_ tetramer; in contrast, median frequency of Rv0125_81-92_ tetramer+ CD4 T cells was relatively low at 0.20% (Figure 4B-E). Comparing the frequency of individual tetramer staining within the same sample, there did not appear to be granulomas that had consistently high frequencies of every tetramer, but instead specific tetramer frequency varied within granulomas (Figure 4E), suggesting that T cells of different antigen specificities were present across granulomas even in the same animal, most easily seen in animal 23820 (gold plot, Figure 4E).

**Figure 4:**
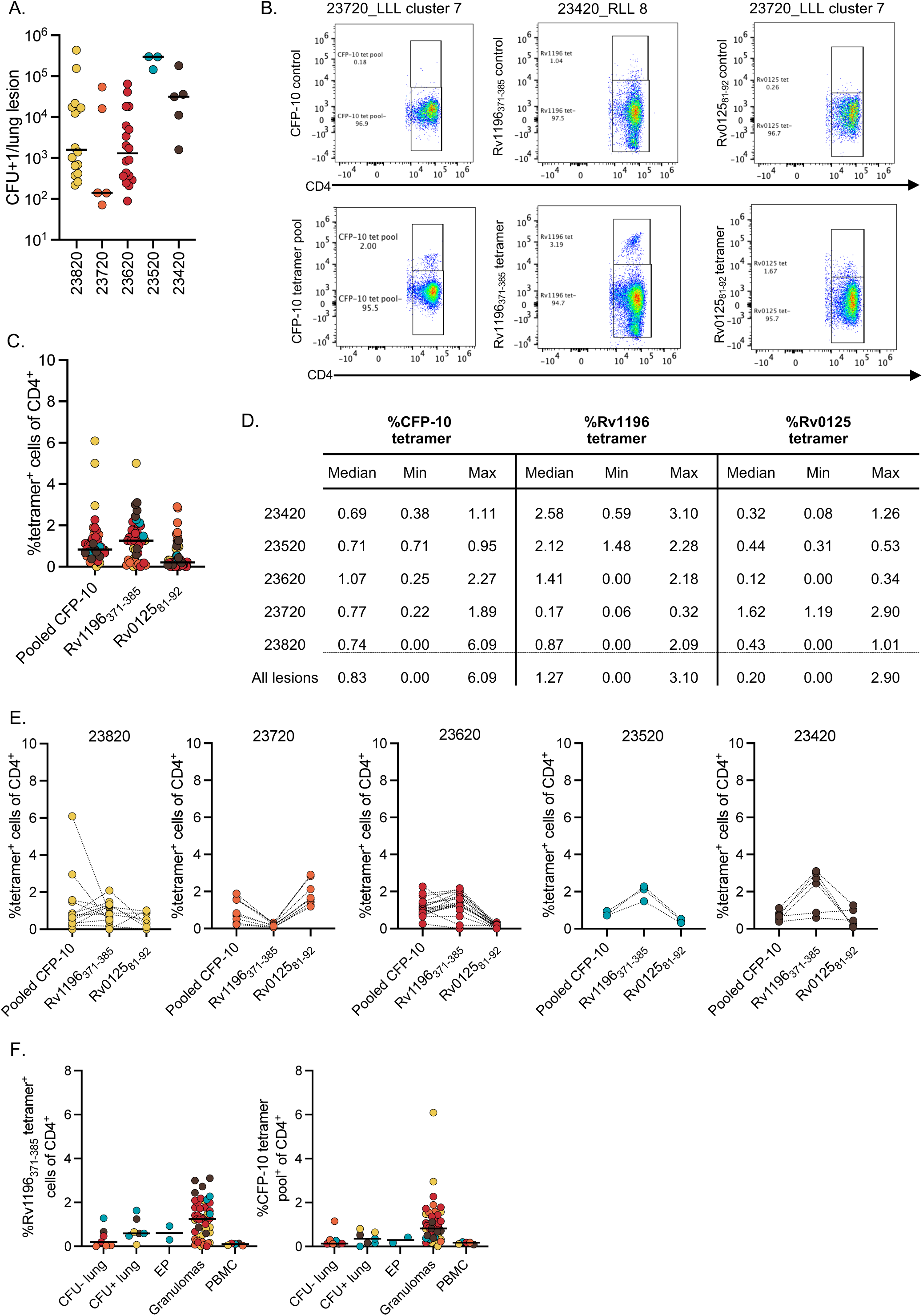
Identification of tetramer^+^ cells in lung lesions. (A) The range in bacterial burden per lung lesion in each macaque, bars represent median values. (B) Representative flow cytometry plots showing the frequency of control tetramers (top) and pooled CFP-10 tetramer^+^ cells, Rv1196_371-385_ tetramer^+^ cells, and Rv012581-92 tetramer^+^ cells (bottom) in CD4^+^ T cells in lung lesions. (C) Frequency of each tetramer+ population of CD4^+^ T cells in lung lesions (individual points) colored by animal. Bars represent median values. (D) Median, minimum and maximum frequencies for CFP-10, Rv1196, and Rv0125 tetramer^+^ cells among all CD4^+^ T cells for each macaque. (E) Frequencies of individual tetramer^+^ cells of CD4^+^ T cells within each animal in which lines connect the same lung lesion sample (animal number stated above each graph). (F) Frequency of Rv1196_371-385_ tetramer^+^ cells and CFP-10 tetramer^+^ cells in CFU^-^ lung samples, CFU^+^ lung samples, extra-pulmonary (EP) granulomas (liver and spleen), lung granulomas and necropsy PBMCs.

As anticipated, when comparing the frequency of tetramer^+^ cells in blood to the site of infection, i.e. lung granulomas, there was an ∼12-fold increase in Rv1196 tetramer^+^ cells in granulomas (median PBMC: 0.1%; median lung granulomas: 1.24% Rv1196) (Figure 4F). There was a 6-fold increase in the frequency of tetramer^+^ cells in lung granulomas as compared to uninvolved lung (CFU=0) (uninvolved lung: 0.19%, median in lung granulomas: 1.24%) or even CFU^+^ lung samples (without obvious granulomas) (Figure 4F). Extrapulmonary granulomas (liver or spleen), although fewer samples were available, also had low frequencies of Rv1196 tetramer^+^ cells (Figure 4F). Similar trends were observed when comparing frequencies of CFP-10 tetramer^+^ cells (Figure 4F). These data support that lung granulomas are enriched sites for tetramer^+^ cells.

### Transcription factor and activation marker expression in tetramer^+^ lung granuloma cells

To assess potential phenotypic and functional differences between tetramer^+^ and tetramer^neg^ CD4 T cells, granuloma samples were stained with the lineage specifying transcription factors T-bet, GATA3, Foxp3, RORγT, and ROR*α*. Although MHC Class I tetramers in other disease models have been used in conjunction with intracellular cytokine staining, few studies have used intracellular cytokine staining in coordination with MHC Class II tetramers (53, 54). One of the primary reasons for this is the instability of the CD4 TCR on the cell surface following TCR ligation (55). Most protocols strongly recommend the use of a protein kinase inhibitor (PKI), such as dasatinib, for stabilizing the TCR to enhance the staining potential of these cells. However, PKIs limit T cell activation and cytokine secretion (56). Thus, we used antibodies specific for transcription factors as a surrogate for cellular functionality along with our Rv1196 and CFP10 tetramers in our flow cytometry panel for analyzing PBMCs, lungs, LNs and lung granulomas (Supplementary Figure 2B-D). For these analyses Rv0125_81-92_ tetramer^+^ cells were excluded due to their relatively low frequencies in these samples.

Significantly higher frequencies of T-bet and RORγT expression were observed for Rv1196_371-385_ and CFP-10 tetramer^+^ cells when compared to tetramer^neg^ cells in the same sample (Figure 5A-B). Frequencies for the remaining transcription factors were low in all CD4^+^ cells; despite this, there was an overall trend of higher expression of GATA3 in tetramer^neg^ cells compared to tetramer^+^ cells. There was a significantly higher frequency of ROR*α* expressing CFP-10 tetramer^+^ cells compared to CFP-10 tetramer^neg^ cells (Figure 5B). There were no observable differences in the frequency of Foxp3 expression between tetramer^+^ and tetramer^neg^ cells; taken together with its low overall frequency indicates that it is not highly expressed in CD4 T cells in MCM lung granulomas at the time points assessed (Figure 5A-B).

**Figure 5:**
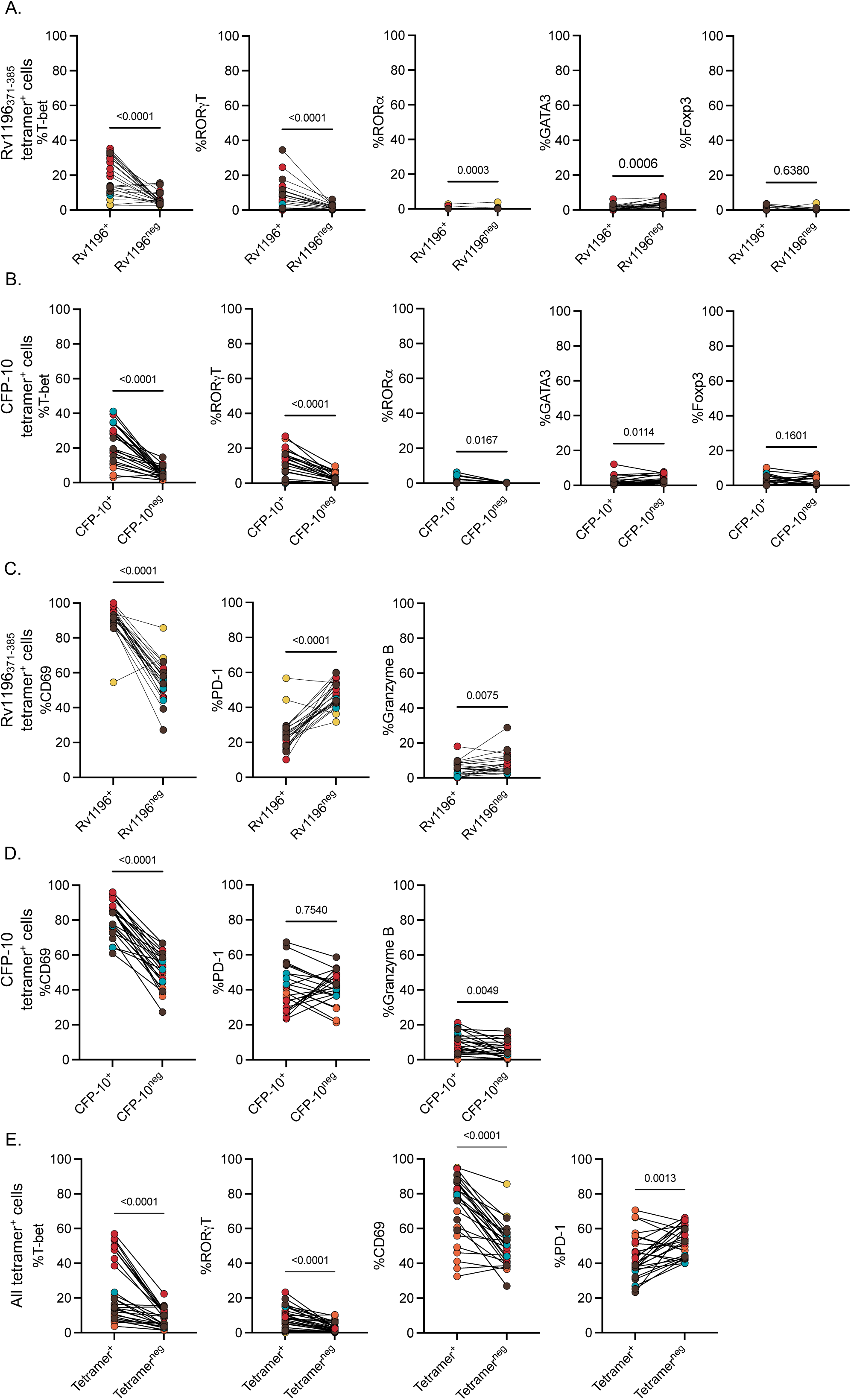
Transcription factor and activation marker expression in tetramer^+^ granuloma CD4^+^ T cells. (A) Frequency of transcription factor expression in Rv1196_371-385_ tetramer^+^ CD4 T cells compared to Rv1196_371-385_ tetramer^neg^ CD4 T cells within the same granuloma sample. (B) Frequency of transcription factor expression in CFP-10 tetramer^+^ cells compared to CFP-10 tetramer^neg^ cells within the same granuloma sample. (C, D) Frequency of activation marker expression (CD69 and PD-1) and granzyme B in Rv1196_371-385_ tetramer^+^ CD4 T cells compared to Rv1196_371-385_ tetramer^neg^ CD4 T cells (C) and CFP-10 tetramer^+^ CD4 T cells compared to CFP-10 pooled tetramer^neg^ CD4 T cells (D) in granulomas. (E) Frequency of T-bet, RORγT, CD69, and PD-1 expression in all tetramer^+^ compared to tetramer^neg^ cells using Boolean OR gating for all 4 tetramers. For all plots each dot represents an individual granuloma or pooled granulomas colored by animal (color legend in Figure 2) with lines connecting the same sample for tetramer+ vs tetramer^neg^ CD4 T cells. Statistical analyses were performed using a Wilcoxon matched-pairs signed rank test.

We evaluated tetramer^+^ cells for activation marker expression (CD69 and PD-1) and production of the cytolytic molecule granzyme B. CFP-10 tetramer^+^ cells and Rv1196_371-385_ tetramer^+^ cells had significantly higher expression of CD69 when compared to tetramer^neg^ cells in the same sample, indicating that Mtb specific CD4 T cells have an activated phenotype (Figure 5C-D). When evaluated for PD-1, Rv1196_371-385_ tetramer^+^ cells had significantly lower frequencies of PD-1 compared to tetramer^neg^ cells (Figure 5C); this trend was not uniform when analyzing CFP-10 tetramer^+^ and tetramer^neg^ cells but instead was animal dependent (Figure 5D). Expression of granzyme B varied based on sample and specific tetramer, with significantly higher frequencies of granzyme B expression in CFP10 tetramer+ cells, and the opposite for Rv1196 tetramer+ cells (higher frequency in tetramer^neg^ cells), suggesting that each group of tetramer^+^ cells may differ in effector function (Figure 5C-D). Despite variability in animal and tetramer, some tetramer^+^ cells express moderate amounts of granzyme B with ranges between 0.0-18.0% for Rv1196_371-385_ tetramer^+^ cells and 0.0-21.2% for CFP-10 tetramer^+^ cells.

There were congruent results when using Boolean gating or individual gating, with higher frequencies of T-bet and RORγT in all tetramer^+^ cells than in tetramer^neg^ cells in the same sample (Figure 5E). Similarly, when CD69 and PD-1 expression was analyzed on all tetramer^+^ CD4 T cells, there were significantly higher frequencies of CD69 and lower frequencies of PD-1 compared to tetramer^neg^ CD4 T cells within the same sample (Figure 5E).

### Transcription factor expression in tetramer^+^ cells negatively correlates with individual granuloma bacterial burden

To investigate the potential relationship between tetramer frequency and bacterial burden within individual lesions, we compared the log_10_ CFU with the frequency of Rv1196 tetramer^+^ or CFP-10 tetramer^+^ cells on an individual granuloma basis (i.e. only including individual isolated granulomas (Figure 6A). There was a significant positive correlation (r = 0.4374, p = 0.0034) between Rv1196 tetramer^+^ cells and CFU and no correlation between CFP-10 tetramer^+^ cells and CFU (r = -0.0164, p = 0.9169) (Figure 6A). In contrast, when function (transcription factor expression) was included in the analysis, there was a significant negative correlation between CFU and the frequency of T-bet (r = -0.5625, p = 0.0122) or RORγT (r = -0.5187, p = 0.0229) expression within Rv1196 tetramer^+^ cells (Figure 6B). This significant negative correlation was also observed between CFU and the expression of RORγT within CFP-10 tetramer^+^ cells (r = -0.5126, p = 0.0354) but not expression of T-bet (r = 0.1080, p = 0.6880) (Figure 6C). Thus, bacterial burden may drive increased numbers of Mtb-specific CD4 T cells in granulomas, but only functional Mtb-specific CD4 T cells are associated with reduced bacterial burden.

**Figure 6:**
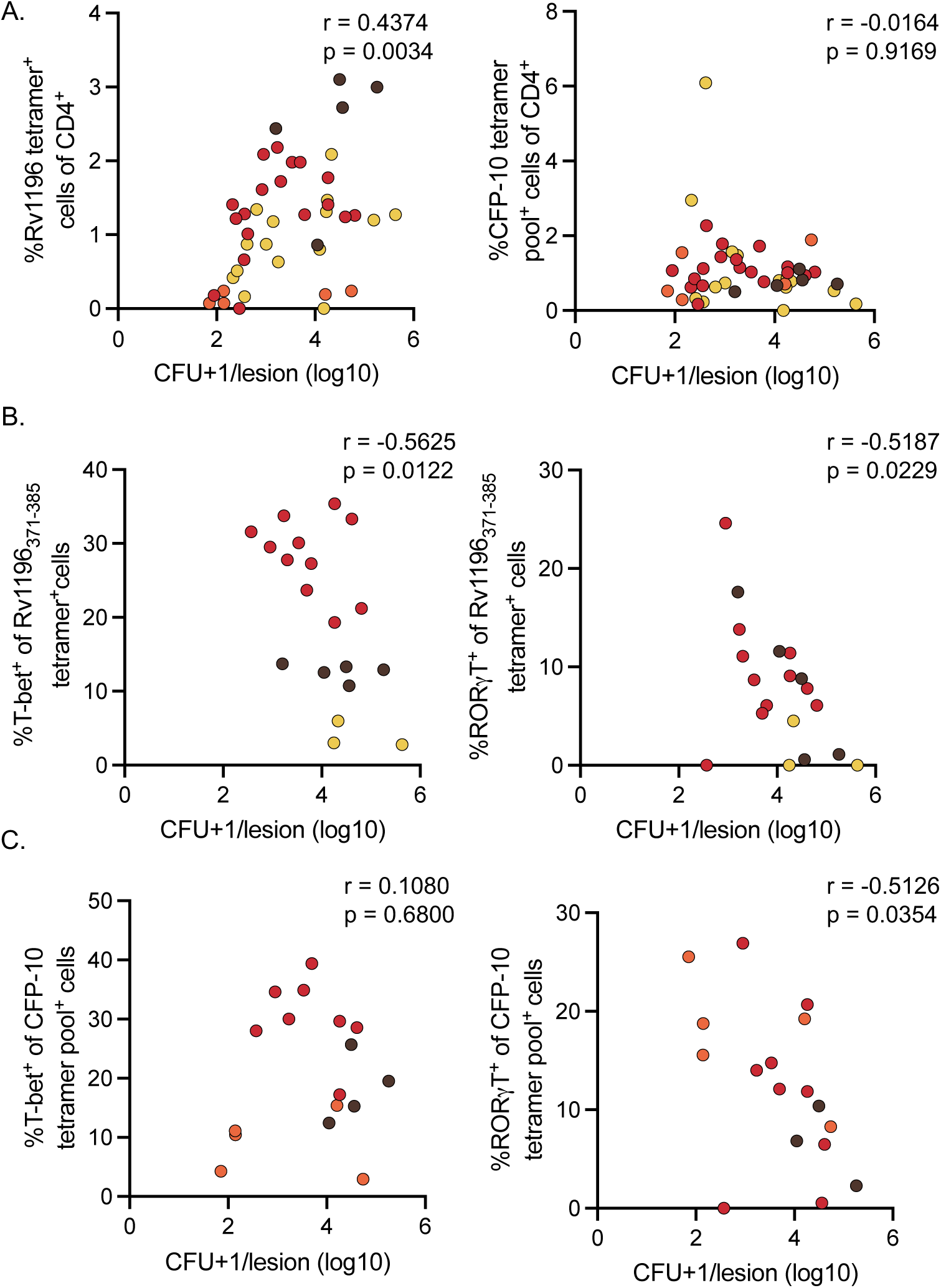
Transcription factor expression in tetramer^+^ cells negatively correlates with individual granuloma bacterial burden. (A) Significant positive correlation between the frequency of Rv1196 tetramer^+^ cells of CD4^+^ T cells and log_10_ transformed CFU+1/lesion (left). No correlation between the frequency of CFP-10 tetramer+ cells of CD4^+^ T cells and log_10_ transformed CFU+1/granuloma (right, Spearman’s r reported for non-parametric). (B) Significant negative correlations between the frequency of T-bet (left) and RORγT (right) expression between Rv1196 tetramer^+^ CD4 T cells and log_10_ transformed CFU+1/granuloma. (C) No correlation between the frequency of T-bet expression among CFP-10 pooled tetramer^+^ cells and log_10_ transformed CFU+1/granuloma (left). Significant negative correlation between the frequency of RORγT expression among CFP-10 tetramer+ cells and log_10_ transformed CFU+1/granuloma (right). Unless otherwise noted, Pearson’s r reported.

## DISCUSSION

Identifying Mtb-specific T cells in granulomas is critical to understanding the functional capabilities of these cells. Unlike viruses, Mtb expresses thousands of antigens and studies in Mtb-infected humans demonstrate that T cells can recognize a wide range of these antigens with substantial variability among people (57). The full range of antigens recognized by T cells in granulomas is unknown. It is also likely that there are many T cells in granulomas that are not specific for Mtb but instead migrate to the granuloma due to inflammatory signals. Limiting analysis to Mtb specific T cells thus provides a focused interrogation of the function of these cells within granulomas. In this study, we developed tetramers that identify T cells specific for two Mtb antigens, Rv1196 and Rv0125, and used these in addition to two previously published CFP-10 tetramers (another Mtb antigen) to study the enrichment and function of these cells in lung granulomas from Mtb-infected macaques. Data from our lab over the past two decades support that most Mtb-infected macaques have T cells that recognize these antigens. Although tetramers specific for Mtb antigens have been developed and used in mice for several years, mice do not form human-like granulomas and the disease trajectory is dissimilar to humans. In contrast, NHP models have many similarities to human TB, including granuloma structure, other pathologies and a wide range of Mtb infection outcomes. However, the rhesus and Chinese cynomolgus macaque MHC loci are extremely complex, even more so than the MHC in humans (35). Thus, we took advantage of the MCM model which has relatively restricted and well characterized MHC alleles for our tetramer development and testing (29, 58, 59). Our data support that granulomas are enriched for Mtb-specific CD4 T cells which have an activated Th1 or Th17 phenotype. Mtb specific CD4 T cells expressing key transcription factors T-bet or RORγT were associated with lower bacterial burden in granulomas, supporting that these cells are performing functions critical to bacterial control.

MHC Class II tetramers for the Mtb protein CFP-10 were described by collaborators (3) and were available through the NIH Tetramer Core. For Mtb antigens Rv1196 and Rv0125, we identified the dominant epitope and MHC allele restriction. Rv1196 is a member of the PPE gene family which is conserved across Mtb strains and *M. bovis* but is absent in other mycobacterial species (8). Though the function remains unknown, Rv1196 has been shown to induce IFN-γ responses and T cell proliferation in PBMCs from human PPD^+^ patients lacking evidence of TB disease, suggesting that it elicits a protective response (8). The second protein investigated in this study, Rv0125, also elicits IFN-γ responses in PPD^+^ patients without evidence of TB disease (7). A predicted secreted serine protease, Rv0125 is conserved across species within the Mtb complex and *M. bovis*, but not in environmental mycobacteria (7). Given their conserved nature and ability to elicit IFN-γ responses within PPD^+^ protected individuals, Rv1196 and Rv0125 were included in the adjuvanted vaccine M72F which has been shown to protect against development of TB disease in subjects with asymptomatic Mtb infection and as a boost to the BCG vaccine in mice and rabbits (9, 60–64).

Here we identified that the dominant epitope from both proteins was restricted by the MHC Class II *Mafa*-DPA1*07:02/*Mafa*-DPB1*19:03 alleles which present antigens to CD4 T cells. The dominant epitope binding region, a.a. 371-385 in Rv1196, contained 2 amino acids, valine and methionine (a.a. 383 and 384) necessary for eliciting an IFN-γ response. We did not observe this phenomenon with Rv0125 but determined that the optimal binding region is smaller than that of Rv1196. Using tetramers designed for these proteins and for the previously mapped CFP-10, we identified Mtb-specific CD4 T cells in various compartments including the airways, uninvolved lung lobes, peripheral blood, thoracic LNs and lung granulomas at necropsy. Comparing the frequencies of tetramer^+^ cells in uninvolved lung or the periphery with the frequencies observed in lung granulomas revealed that granulomas are enriched for Mtb-specific (tetramer^+^) CD4 T cells.

The phenotype of Mtb-specific CD4 T cells in NHP TB lung granulomas using tetramer staining had not been previously explored. Here we set out to identify a functional phenotype for tetramer^+^ cells using flow cytometry by staining for transcription factors, activation markers, and a cytolytic effector molecule. Two transcription factors associated with T cell control of Mtb include T-bet and RORγT (51, 65, 66). T-bet is a transcription factor that induces several effector molecules, including production of pro-inflammatory cytokines, cytotoxic effectors, chemokines, and regulation of T cell responses (67). In some studies, particularly in the context of vaccination, RORγT and the Th17 phenotype has been associated with protection against or control of Mtb infection (65, 66, 68). Rv1196_371-385_ tetramer^+^ cells and CFP-10 tetramer^+^ CD4 T cells in granulomas had significantly higher frequencies of T-bet and RORγT expression compared to tetramer^neg^ cells. ROR*α*, GATA3, and Foxp3 expression was relatively low in tetramer^+^ cells compared to T-bet or RORγT expression. These data reveal that Rv1196 and CFP-10 tetramer^+^ cells exhibit a Th1 and/or Th17 phenotype. However, in our and other’s previous studies of NHP granulomas, either by flow cytometry or single cell RNA sequencing, only very low frequencies of IL-17^+^ T cells were observed (20, 21, 69, 70); we surmised previously that the RORγT^+^ T cells might be ex-Th17 cells (69, 71).

There were significantly higher frequencies of CD69 expression on tetramer^+^ CD4 T cells compared to tetramer^neg^ CD4 T cells within the same granuloma. CD69 is a cell surface marker upregulated following T cell activation, though it also can play a role in cytokine release and cellular migration (72, 73). In contrast, PD-1 expression, which is both a marker for T cell activation or chronically stimulated cells, was significantly lower in Rv1196_371-385_ tetramer^+^ cells as compared to tetramer^neg^ cells, although it varied within animal for CFP-10 tetramer^+^ cells. This dynamic expression amongst tetramer^+^ cells of higher CD69 and lower PD-1 may indicate that these cells were not recently stimulated, but instead express CD69 as a function of immune environment and for its contribution to other effector functions. Though previous studies have shown low levels of exhaustion marker expression in T cells within NHP and human lung granulomas, this is the first direct evidence of low levels of PD-1 expression on Mtb-specific T cells in a macaque model (21, 23). Since granzyme B expression is more commonly associated with CD8 cytotoxic T cells, we expected low levels in the CD4 tetramer^+^ cells. However, since our gating included CD4^+^CD8^+^ T cells, we investigated the presence of granzyme B within tetramer^+^ cells and tetramer^neg^ cells, finding significantly higher levels of granzyme B expression within CFP-10 tetramer^+^ cells and lower frequencies in Rv1196 tetramer^+^ cells, suggesting that Mtb-specific cells for different antigens may have different functional capacities.

By comparing all tetramer^+^ cells to tetramer^neg^ cells in individual granulomas we observed higher frequencies of T-bet, RORγT, and CD69 and significantly lower levels of PD-1 in tetramer+ cells. This reinforces that Mtb-specific T cells are activated and better poised for functionality in granulomas than Mtb-nonspecific CD4 T cells. Although we only captured a small fraction of Mtb-specific CD4 T cells due to the large antigenic repertoire of mycobacteria, these data suggest that granulomas contain a population of Mtb-non-specific CD4 T cells that migrate to the site of infection but are unlikely to participate in control of infection.

The variable frequency of tetramer^+^ T cells observed in lung granulomas within the same animal and between animals emphasizes the independent nature of granulomas. In addition, our data suggest that recruitment of Mtb-specific cells is not solely based on optimal T cell recruiting granuloma phenotypes. For instance, we did not observe granulomas with higher frequencies of Rv1196_371-385_ tetramer^+^ cells also having higher frequencies of CFP-10 tetramer^+^ cells as might be predicted, with the idea that some granulomas may have a better capacity for Mtb-specific T cell recruitment through the production of chemokines or other cell signals. Rather, the data presented here suggest that T cells with different Mtb antigen specificities vary across granulomas even within the same animal. One potential hypothesis for this observation is that the first Mtb-specific T cells recruited to a granuloma become the dominant specific cells through local proliferation. The levels of tetramer^+^ cells within granulomas also likely depends on the state of Mtb within those granulomas (i.e. quiescent or replicating, or high levels of dead bacteria). We observed a significant positive correlation between the frequency of Rv1196 tetramer^+^ cells and CFU in individual lesions. This suggests that as bacterial burden increases, so does recruitment or replication of specific T cells within individual lesions. However, we observed a significant negative correlation between the frequency of T-bet or RORγT within tetramer^+^ cells, suggesting the functionality of Mtb-specific cells is critical for reducing bacterial burden.

Although the initial screening for epitopes was expected to identify both MHC Class I and MHC Class II epitopes, we only identified those recognized by CD4 T cells for both Rv1196 and Rv0125 proteins. This likely represents a limitation of our system, particularly in the use of IFN-γ as a functional readout as CD8 T cells from NHP often express low levels of IFN-γ(20). In addition, CD4 T cells may outgrow the CD8 T cells in stimulated PBMC cultures. In the future, mapping peptides that elicit CD8 T cell responses could be achieved by modifying and optimizing ELISPOT protocols for cytolytic molecules and peptide pools with 8-12 amino acids in length and by depleting CD4 T cells prior to T cell culture. We identified limited frequencies of Rv0125 tetramer^+^ cells and to some degree CFP-10 tetramer^+^ cells in this study. Two potential hypotheses for this may be that Rv0125 is not highly produced by Mtb in vivo or that the binding affinity needed for identification of tetramer^+^ cells is higher than that needed to elicit IFN-γproduction. We also consistently observed low IFN-γ responses to the Rv0125 peptide pool in stimulated PBMCs, suggesting that this protein may contain other masking antigens, i.e. those that preferentially bind to MHC II molecules but do not elicit IFN-γ responses. One limitation to the use of tetramers in the context of a large bacterial pathogen (i.e. many proteins produced) is that it provides a narrow view regarding the function of a small proportion of the potential specific cells. However, IFN-γ ELISPOTS performed using pools of 54 and 300 immunodominant peptides elicit similar levels of IFN-γresponses in NHPs, suggesting that there is a smaller set of Mtb proteins responsible for the majority of the IFN-γMtb response (2).

A final limitation of these studies was the inability to stain for cytokines in conjunction with tetramers, given the need for inclusion of dasatinib in the tetramer staining protocol to stabilize the MHC Class II tetramer binding. Although we gained substantial information by including transcription factor staining, we were unable to assess the true effector functions of these Mtb-specific T cells. However, now that the tetramers are available, more advanced techniques such as CITE-Seq may allow future transcriptional analysis of the functions of Mtb specific cells, representing an additional advance in our understanding of T cell responses in granulomas.

In summary, our data demonstrate that granulomas are enriched sites for Mtb-specific CD4 T cells as compared to lung tissue, LNs, or blood. While we can identify tetramer^+^ cells in the airways as early as 3 weeks post infection, the frequencies of cells are low and do not increase until 8 weeks post infection. Within lung granulomas, our data revealed that the majority of Mtb-specific CD4 T cells are CD69^+^ Th1 cells or Th17 (or ex-Th17) cells. We demonstrated a significant negative correlation between expression of T-bet or RORγT within tetramer^+^ cells and bacterial burden in granulomas, highlighting the importance of functional Mtb-specific T cells in reducing bacterial burden within lung granulomas. The use of tetramers provides insight into the phenotype of granuloma CD4 Mtb-specific cells, suggesting that enhancing or inducing Th1/Th17 functionality in Mtb-specific CD4 T cells would be advantageous in the context of vaccine development.

## AUTHOR CONTRIBUTIONS

Conceptualization, JLF and NLG; investigation, NLG, KK, PM, PLL, and HA; resources, JLF, CS, and SO; formal analysis, NLG, PM, and JLF; writing-original draft, NLG and JLF; writing-reviewing and editing, NLG, PM, PLL, JLF; supervision, CS and JLF; funding acquisition, JLF.

## ACKNOWLEDGEMENTS

This work was supported by NIH R01 AI123093 (JLF), NIH NIAID 75N93019C00071 (JLF), NIH R01 AI111815 (CS & SLO), T32 AI065380 (NLG), and the Bill and Melinda Gates Foundation (JLF). We are grateful to and appreciative of the lab members of the Flynn, Scanga, Lin, and Mattila labs, in particular Ryan Kelly, Jennie Vorhauer, Kush Patel, Abigail Gubernat, Cassaundra Ameel, Mark Rodgers, and Janelle Gleim for their dedication in preparing the necropsy samples from these animals. In addition, we thank our veterinary technicians Jaime Tomko, Dan Fillmore, Kara Kracinovsky, and Jennifer Sakal for their exceptional care of the animals and for performing all procedures, and L. James Frye and Jaime Tomko for performing PET CT imaging. We are grateful to Amy Ellis-Connell and Alexis Balgeman for providing reagents and guidance on epitope mapping. Lastly, we gratefully acknowledge the work of the NIH Tetramer Facility for providing valuable resources.

## Tetramer study figure legends

**Supplementary figure 1:**
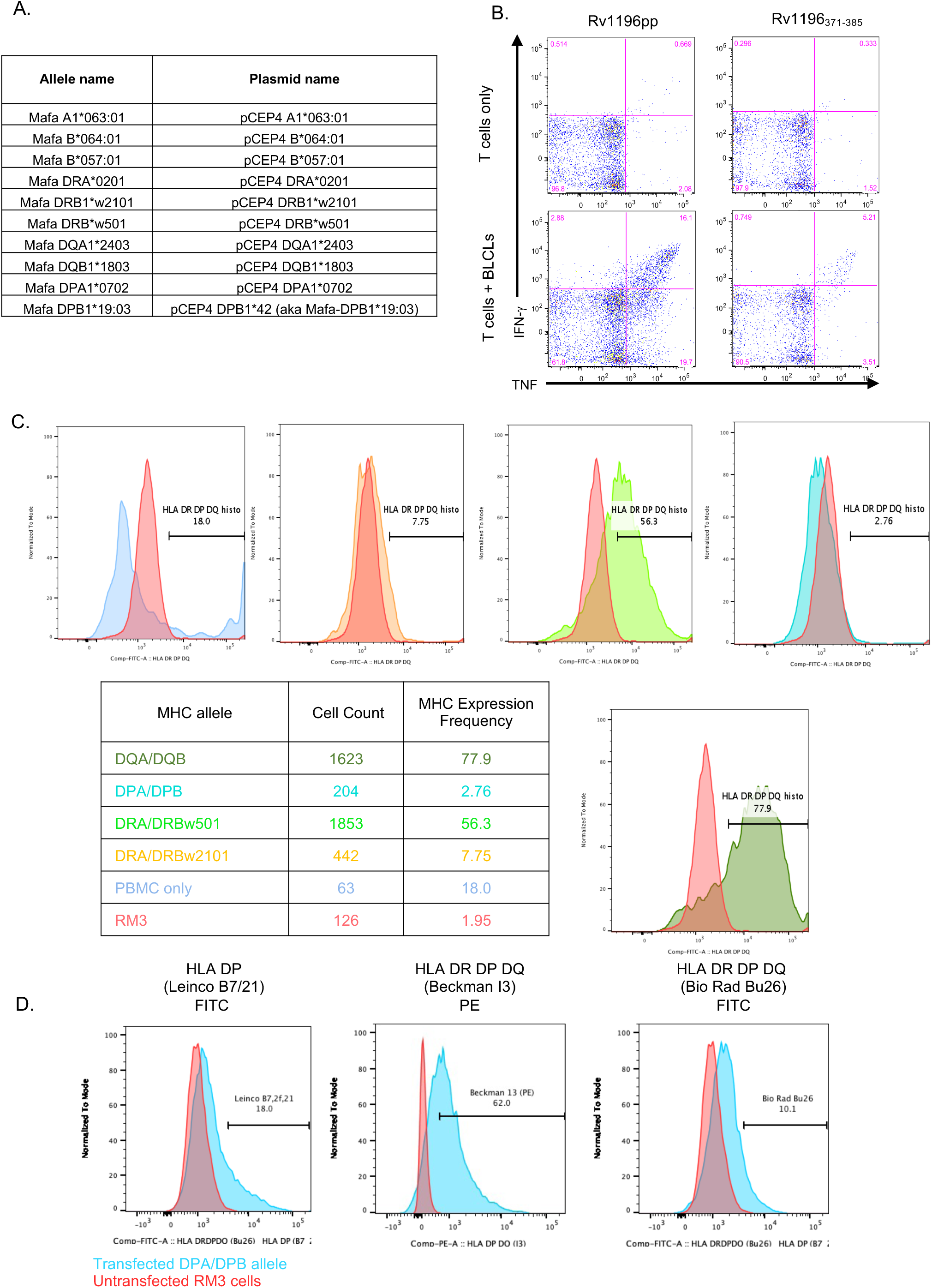
T cell culture staining, plasmids, and RM3 cell transfection plots. (A) Table of plasmid reagents for transfection of RM3 cells. (B) Expanded T cells were co-cultured with peptide-pulsed (Rv1196 or Rv1196_371-385_) irradiated BLCLs and flow cytometry performed for pro-inflammatory cytokine responses (IFN-γ and TNF) to determine whether the Rv1196_371-385_ was presented to CD4 or CD8 T cells. Gating strategy was performed using live/dead exclusion followed by gating on singlets, lymphocytes, CD3^+^ and CD4^+^ events. (C) Flow plots showing the detection of DR, DP, and DQ alleles (colored according to table) following RM3 transfection. Limited expression can be observed in RM3 cells transfected with DPA/DPB (teal) and DRA/DRBw*2101 (orange) alleles as compared to untransfected RM3 cells. (D) Testing of three additional antibodies (antibody, clone, and fluorophore listed above each graph) showing the detection of DPA/DPB allele expression in transfected RM3 cells.

**Supplementary figure 2:**
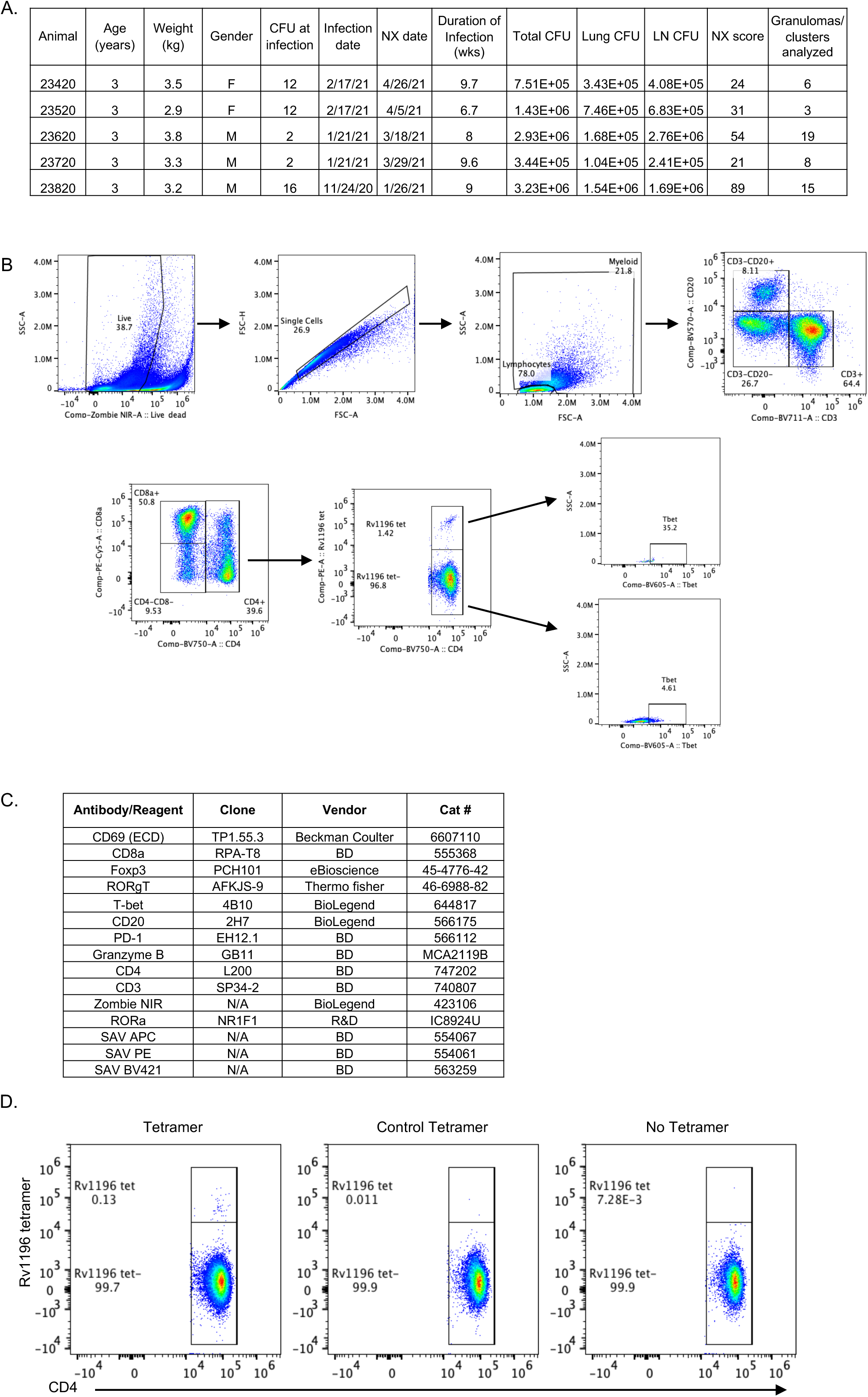
Animals used for tetramer^+^ CD4 T cell identification studies, gating strategy for tissue samples from necropsy, and flow reagents. (A) Table of MCMs used for identification of tetramer^+^ cells in the periphery, BAL, and necropsy samples (B) Flow cytometry gating tree for necropsy samples (example shown is Animal #23620, RLL granuloma cluster 12): Events were gated on live cells using Zombie NIR, single cell events were selected from live events (FSC-A vs FSC-H), and lymphocytes identified from single cells based on FSC-A and SSC-A. Lymphocytes were gated based on CD3 and CD20 expression with T cells (CD3^+^) subsequently gated on CD4^+^ and CD8a^+^ expression. All CD4^+^ cells were gated for tetramer^+^ events (CD4 on x axis and tetramer on y axis). All tetramer^+^ and tetramer^neg^ cells were gated for transcription factors, granzyme B or activation marker expression. (C) Table containing antibodies used for staining necropsy samples for flow cytometry analysis. (D) Example flow cytometry plots showing gating for Rv1196_371-385_ tetramer^+^ cells in tetramer, control tetramer, or no tetramer stained PBMCs at necropsy from animal #23420. Gating strategy was performed using live/dead exclusion followed by gating on singlets, lymphocytes, CD3^+^ and CD4^+^ events as described in (B).

**Supplementary table 1:**
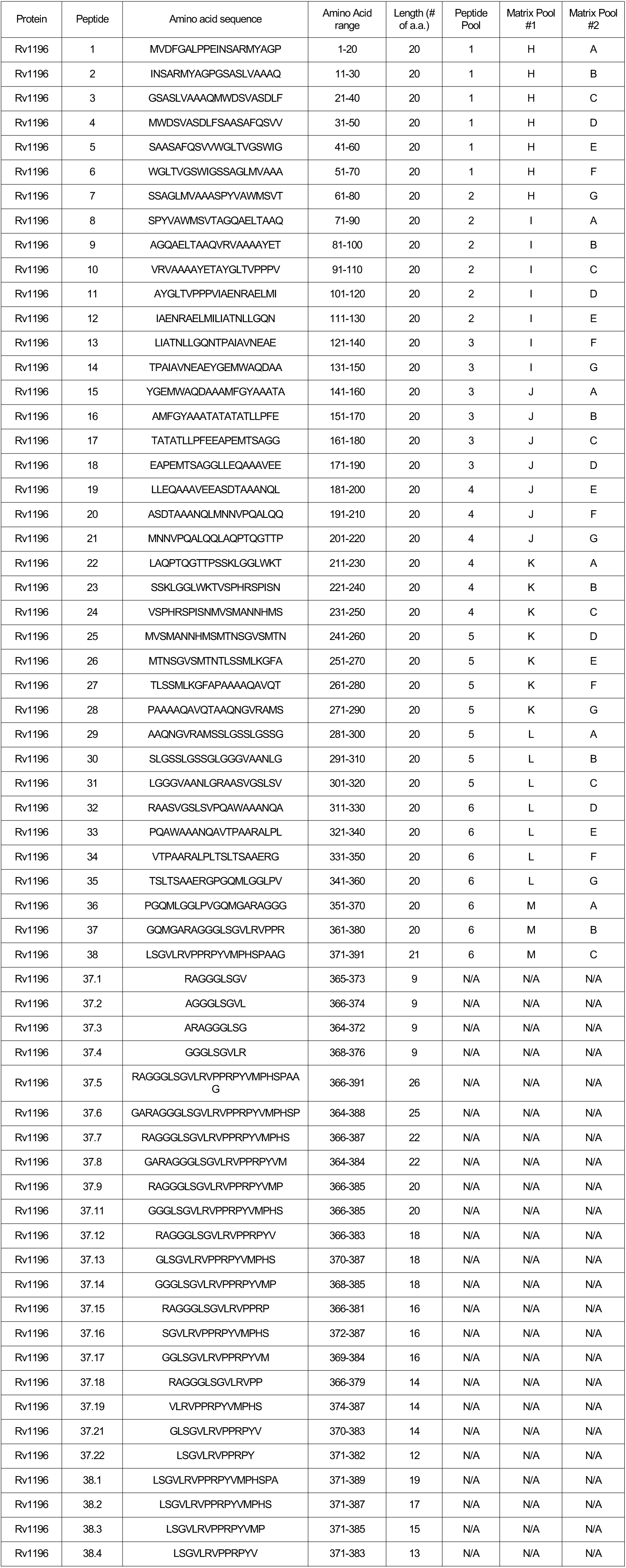
**Peptides used for mapping Rv1196**

**Supplementary table 2:**
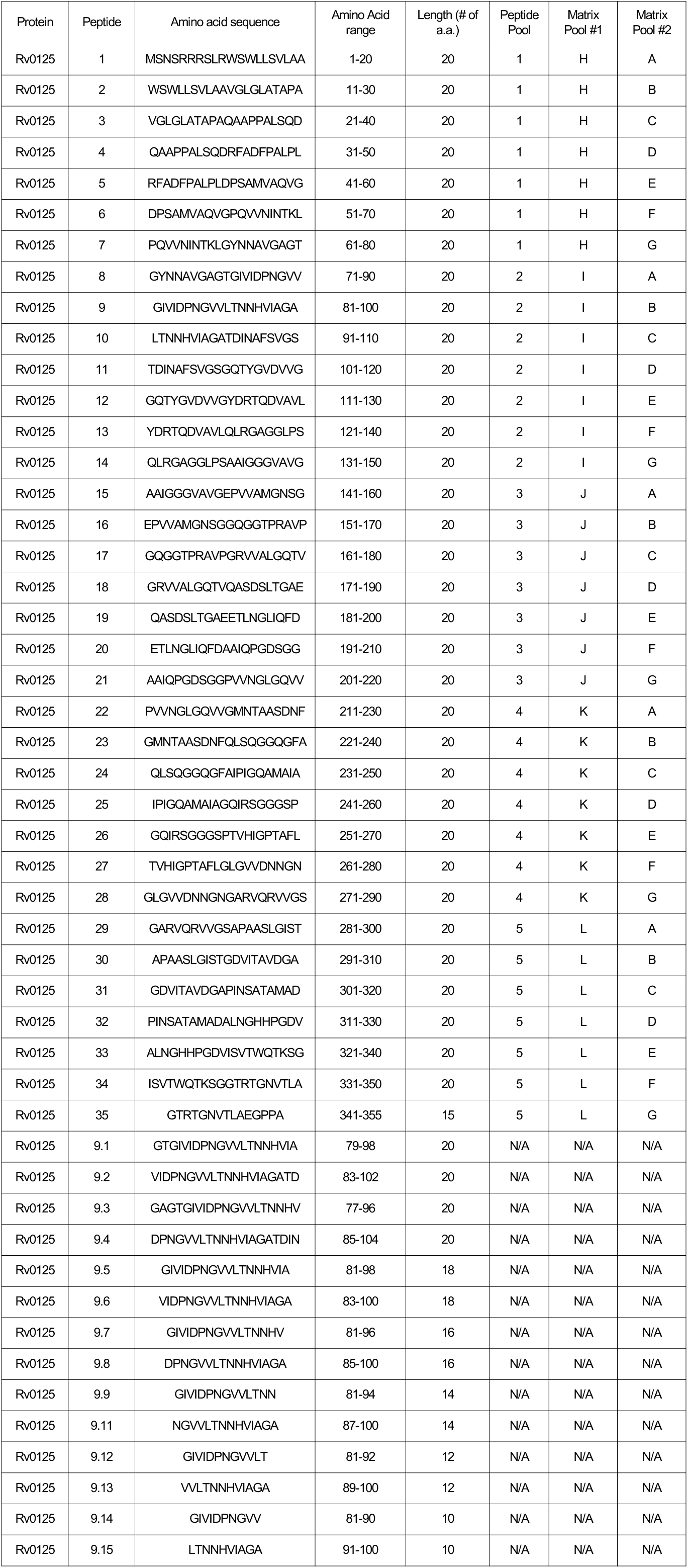
**Peptides used for mapping Rv0125**

## REFERENCES

1. Gomez, M., S. Johnson, and M. L. Gennaro. 2000. Identification of secreted proteins of Mycobacterium tuberculosis by a bioinformatic approach. Infect Immun 68: 2323–2327.

2. Mothé, B. R., C. S. Lindestam Arlehamn, C. Dow, M. B. C. Dillon, R. W. Wiseman, P. Bohn, J. Karl, N. A. Golden, T. Gilpin, T. W. Foreman, M. A. Rodgers, S. Mehra, T. J. Scriba, J. L. Flynn, D. Kaushal, D. H. O’Connor, and A. Sette. 2015. The TB-specific CD4(+) T cell immune repertoire in both cynomolgus and rhesus macaques largely overlap with humans. Tuberculosis (Edinb) 95: 722–735.

3. Ellis, A., A. Balgeman, M. Rodgers, C. Updike, J. Tomko, P. Maiello, C. A. Scanga, and S. L. O’Connor. 2017. Characterization of T Cells Specific for CFP-10 and ESAT-6 in Mycobacterium tuberculosis-Infected Mauritian Cynomolgus Macaques. Infect Immun 85.

4. Ravn, P., A. Demissie, T. Eguale, H. Wondwosson, D. Lein, H. A. Amoudy, A. S. Mustafa, A. K. Jensen, A. Holm, I. Rosenkrands, F. Oftung, J. Olobo, F. von Reyn, and P. Andersen. 1999. Human T cell responses to the ESAT-6 antigen from Mycobacterium tuberculosis. The Journal of infectious diseases 179: 637–645.

5. Arend, S. M., P. Andersen, K. E. van Meijgaarden, R. L. Skjot, Y. W. Subronto, J. T. van Dissel, and T. H. Ottenhoff. 2000. Detection of active tuberculosis infection by T cell responses to early-secreted antigenic target 6-kDa protein and culture filtrate protein 10. The Journal of infectious diseases 181: 1850–1854.

6. Kamath, A. B., J. Woodworth, X. Xiong, C. Taylor, Y. Weng, and S. M. Behar. 2004. Cytolytic CD8+ T cells recognizing CFP10 are recruited to the lung after Mycobacterium tuberculosis infection. J Exp Med 200: 1479–1489.

7. Skeiky, Y. A., M. J. Lodes, J. A. Guderian, R. Mohamath, T. Bement, M. R. Alderson, and S. G. Reed. 1999. Cloning, expression, and immunological evaluation of two putative secreted serine protease antigens of Mycobacterium tuberculosis. Infect Immun 67: 3998–4007.

8. Dillon, D. C., M. R. Alderson, C. H. Day, D. M. Lewinsohn, R. Coler, T. Bement, A. Campos-Neto, Y. A. Skeiky, I. M. Orme, A. Roberts, S. Steen, W. Dalemans, R. Badaro, and S. G. Reed. 1999. Molecular characterization and human T-cell responses to a member of a novel Mycobacterium tuberculosis mtb39 gene family. Infect Immun 67: 2941–2950.

9. Leroux-Roels, I., S. Forgus, F. De Boever, F. Clement, M.-A. Demoitié, P. Mettens, P. Moris, E. Ledent, G. Leroux-Roels, and O. Ofori-Anyinam. 2013. Improved CD4+ T cell responses to Mycobacterium tuberculosis in PPD-negative adults by M72/AS01 as compared to the M72/AS02 and Mtb72F/AS02 tuberculosis candidate vaccine formulations: A randomized trial. Vaccine 31: 2196–2206.

10. Green, A. M., R. DiFazio, and J. L. Flynn. 2013. IFN-γ from CD4 T Cells Is Essential for Host Survival and Enhances CD8 T Cell Function during Mycobacterium tuberculosis Infection. The Journal of Immunology 190: 270–277.

11. Tubo, N. J., and M. K. Jenkins. 2014. CD4+ T Cells: guardians of the phagosome. Clin Microbiol Rev 27: 200–213.

12. Höhn, H., C. Kortsik, I. Zehbe, W. E. Hitzler, K. Kayser, K. Freitag, C. Neukirch, P. Andersen, T. M. Doherty, and M. Maeurer. 2007. MHC class II Tetramer Guided Detection of Mycobacterium tuberculosis-specific CD4+ T Cells in Peripheral Blood from Patients with Pulmonary Tuberculosis. Scandinavian Journal of Immunology 65: 467–478.

13. Bronke, C., N. M. Palmer, G. H. Westerlaken, M. Toebes, G. M. van Schijndel, V. Purwaha, K. E. van Meijgaarden, T. N. Schumacher, D. van Baarle, K. Tesselaar, and A. Geluk. 2005. Direct ex vivo detection of HLA-DR3-restricted cytomegalovirus- and Mycobacterium tuberculosis-specific CD4+ T cells. Hum Immunol 66: 950–961.

14. Wei, H., R. Wang, Z. Yuan, C. Y. Chen, D. Huang, L. Halliday, W. Zhong, G. Zeng, Y. Shen, L. Shen, Y. Wang, and Z. W. Chen. 2009. DR*W201/P65 tetramer visualization of epitope-specific CD4 T-cell during M. tuberculosis infection and its resting memory pool after BCG vaccination. PLoS One 4: e6905–e6905.

15. Lindestam Arlehamn, C. S., A. Gerasimova, F. Mele, R. Henderson, J. Swann, J. A. Greenbaum, Y. Kim, J. Sidney, E. A. James, R. Taplitz, D. M. McKinney, W. W. Kwok, H. Grey, F. Sallusto, B. Peters, and A. Sette. 2013. Memory T cells in latent Mycobacterium tuberculosis infection are directed against three antigenic islands and largely contained in a CXCR3+CCR6+ Th1 subset. PLoS pathogens 9: e1003130–e1003130.

16. Gideon, H. P., T. K. Hughes, M. H. Wadsworth, A. A. Tu, T. M. Gierahn, J. M. Peters, F. F. Hopkins, J.-R. Wei, C. Kummerlowe, N. L. Grant, K. Nargan, J. Phuah, H. J. Borish, P. Maiello, A. G. White, C. G. Winchell, S. K. Nyquist, S. K. C. Ganchua, A. Myers, K. V. Patel, C. L. Ameel, C. T. Cochran, S. Ibrahim, J. A. Tomko, L. J. Frye, J. M. Rosenberg, A. Shih, M. Chao, C. A. Scanga, J. Ordovas-Montanes, B. Berger, J. T. Mattila, R. Madansein, J. C. Love, P. L. Lin, A. Leslie, S. M. Behar, B. Bryson, J. L. Flynn, S. M. Fortune, and A. K. Shalek. 2021. Multimodal profiling of lung granulomas reveals cellular correlates of tuberculosis control. bioRxiv: 2020.2010.2024.352492.

17. Millar, J. A., J. R. Butler, S. Evans, J. T. Mattila, J. J. Linderman, J. L. Flynn, and D. E. Kirschner. 2020. Spatial Organization and Recruitment of Non-Specific T Cells May Limit T Cell-Macrophage Interactions Within Mycobacterium tuberculosis Granulomas. Front Immunol 11: 613638.

18. Flynn, J. L., and J. D. Ernst. 2000. Immune responses in tuberculosis. Current Opinion in Immunology 12: 432–436.

19. Ernst, J. D. 2012. The immunological life cycle of tuberculosis. Nature Reviews Immunology 12: 581–591.

20. Gideon, H. P., J. Phuah, A. J. Myers, B. D. Bryson, M. A. Rodgers, M. T. Coleman, P. Maiello, T. Rutledge, S. Marino, S. M. Fortune, D. E. Kirschner, P. L. Lin, and J. L. Flynn. 2015. Variability in tuberculosis granuloma T cell responses exists, but a balance of pro- and anti-inflammatory cytokines is associated with sterilization. PLoS pathogens 11: e1004603.

21. Wong, E. A., L. Joslyn, N. L. Grant, E. Klein, P. L. Lin, D. E. Kirschner, and J. L. Flynn. 2018. Low Levels of T Cell Exhaustion in Tuberculous Lung Granulomas. Infect Immun 86.

22. Kauffman, K. D., M. A. Sallin, S. Sakai, O. Kamenyeva, J. Kabat, D. Weiner, M. Sutphin, D. Schimel, L. Via, C. E. Barry, T. Wilder-Kofie, I. Moore, R. Moore, and D. L. Barber. 2018. Defective positioning in granulomas but not lung-homing limits CD4 T-cell interactions with Mycobacterium tuberculosis-infected macrophages in rhesus macaques. Mucosal Immunology 11: 462–473.

23. McCaffrey, E. F., M. Donato, L. Keren, Z. Chen, A. Delmastro, M. B. Fitzpatrick, S. Gupta, N. F. Greenwald, A. Baranski, W. Graf, R. Kumar, M. Bosse, C. C. Fullaway, P. K. Ramdial, E. Forgó, V. Jojic, D. Van Valen, S. Mehra, S. A. Khader, S. C. Bendall, M. van de Rijn, D. Kalman, D. Kaushal, R. L. Hunter, N. Banaei, A. J. C. Steyn, P. Khatri, and M. Angelo. 2022. The immunoregulatory landscape of human tuberculosis granulomas. Nat Immunol 23: 318–329.

24. Altman, J. D., P. A. Moss, P. J. Goulder, D. H. Barouch, M. G. McHeyzer-Williams, J. I. Bell, A. J. McMichael, and M. M. Davis. 1996. Phenotypic analysis of antigen-specific T lymphocytes. Science 274: 94–96.

25. D. Otting, N., C. M. Heijmans, R. C. Noort, N. G. de Groot, G. G. Doxiadis, J. J. van Rood, D. I. Watkins, and R. E. Bontrop. 2005. Unparalleled complexity of the MHC class I region in rhesus macaques. Proc Natl Acad Sci U S A 102: 1626–1631.

26. Daza-Vamenta, R., G. Glusman, L. Rowen, B. Guthrie, and D. E. Geraghty. 2004. Genetic divergence of the rhesus macaque major histocompatibility complex. Genome Res 14: 1501–1515.

27. Kulski, J. K., T. Anzai, T. Shiina, and H. Inoko. 2004. Rhesus macaque class I duplicon structures, organization, and evolution within the alpha block of the major histocompatibility complex. Mol Biol Evol 21: 2079–2091.

28. E. Watanabe, A., T. Shiina, S. Shimizu, K. Hosomichi, K. Yanagiya, Y. F. Kita, T. Kimura, E. Soeda, R. Torii, K. Ogasawara, J. K. Kulski, and H. Inoko. 2007. A BAC-based contig map of the cynomolgus macaque (Macaca fascicularis) major histocompatibility complex genomic region. Genomics 89: 402–412.

29. Wiseman, R. W., J. A. Karl, B. N. Bimber, C. E. O’Leary, S. M. Lank, J. J. Tuscher, A. M. Detmer, P. Bouffard, N. Levenkova, C. L. Turcotte, E. Szekeres, Jr., C. Wright, T. Harkins, and D. H. O’Connor. 2009. Major histocompatibility complex genotyping with massively parallel pyrosequencing. Nature medicine 15: 1322–1326.

30. Bontrop, R. E. 2006. Comparative genetics of MHC polymorphisms in different primate species: duplications and deletions. Hum Immunol 67: 388–397.

31. Horton, R., R. Gibson, P. Coggill, M. Miretti, R. J. Allcock, J. Almeida, S. Forbes, J. G. R. Gilbert, K. Halls, J. L. Harrow, E. Hart, K. Howe, D. K. Jackson, S. Palmer, A. N. Roberts, S. Sims, C. A. Stewart, J. A. Traherne, S. Trevanion, L. Wilming, J. Rogers, P. J. de Jong, J. F. Elliott, S. Sawcer, J. A. Todd, J. Trowsdale, and S. Beck. 2008. Variation analysis and gene annotation of eight MHC haplotypes: the MHC Haplotype Project. Immunogenetics 60: 1–18.

32. Parham, P. 2005. MHC class I molecules and kirs in human history, health and survival. Nature Reviews Immunology 5: 201–214.

33. Trowsdale, J., and J. C. Knight. 2013. Major Histocompatibility Complex Genomics and Human Disease. Annual Review of Genomics and Human Genetics 14: 301–323.

34. Krebs, K. C., Z. Jin, R. Rudersdorf, A. L. Hughes, and D. H. O’Connor. 2005. Unusually High Frequency MHC Class I Alleles in Mauritian Origin Cynomolgus Macaques. The Journal of Immunology 175: 5230.

35. Wiseman, R. W., J. A. Karl, P. S. Bohn, F. A. Nimityongskul, G. J. Starrett, and D. H. O’Connor. 2013. Haplessly hoping: macaque major histocompatibility complex made easy. Ilar j 54: 196–210.

36. Maiello, P., R. M. DiFazio, A. M. Cadena, M. A. Rodgers, P. L. Lin, C. A. Scanga, and J. L. Flynn. 2017. Rhesus macaques are more susceptible to progressive tuberculosis than cynomolgus macaques: A quantitative comparison. Infection and Immunity.

37. Rodgers, M. A., C. Ameel, A. L. Ellis-Connell, A. J. Balgeman, P. Maiello, G. L. Barry, T. C. Friedrich, E. Klein, S. L. O’Connor, and C. A. Scanga. 2018. Preexisting Simian Immunodeficiency Virus Infection Increases Susceptibility to Tuberculosis in Mauritian Cynomolgus Macaques. Infect Immun 86.

38. Sharpe, S. A., A. D. White, L. Sibley, F. Gleeson, G. A. Hall, R. J. Basaraba, A. McIntyre, S. O. Clark, K. Gooch, P. D. Marsh, A. Williams, and M. J. Dennis. 2017. An aerosol challenge model of tuberculosis in Mauritian cynomolgus macaques. PLoS One 12: e0171906.

39. Lin, P. L., M. Rodgers, L. Smith, M. Bigbee, A. Myers, C. Bigbee, I. Chiosea, S. V. Capuano, C. Fuhrman, E. Klein, and J. L. Flynn. 2009. Quantitative comparison of active and latent tuberculosis in the cynomolgus macaque model. Infect Immun 77: 4631–4642.

40. Capuano, S. V., 3rd, D. A. Croix, S. Pawar, A. Zinovik, A. Myers, P. L. Lin, S. Bissel, C. Fuhrman, E. Klein, and J. L. Flynn. 2003. Experimental Mycobacterium tuberculosis infection of cynomolgus macaques closely resembles the various manifestations of human M. tuberculosis infection. Infection and immunity 71: 5831–5844.

41. White, A. G., P. Maiello, M. T. Coleman, J. A. Tomko, L. J. Frye, C. A. Scanga, P. L. Lin, and J. L. Flynn. 2017. Analysis of 18FDG PET/CT Imaging as a Tool for Studying Mycobacterium tuberculosis Infection and Treatment in Non-human Primates. J Vis Exp: 56375.

42. Calman, A. F., and B. M. Peterlin. 1987. Mutant human B cell lines deficient in class II major histocompatibility complex transcription. J Immunol 139: 2489–2495.

43. Giraldo-Vela, J. P., R. Rudersdorf, C. Chung, Y. Qi, L. T. Wallace, B. Bimber, G. J. Borchardt, D. L. Fisk, C. E. Glidden, J. T. Loffredo, S. M. Piaskowski, J. R. Furlott, J. P. Morales-Martinez, N. A. Wilson, W. M. Rehrauer, J. D. Lifson, M. Carrington, and D. I. Watkins. 2008. The major histocompatibility complex class II alleles Mamu-DRB1*1003 and -DRB1*0306 are enriched in a cohort of simian immunodeficiency virus-infected rhesus macaque elite controllers. J Virol 82: 859–870.

44. Garson, D., M. C. Dokhélar, H. Wakasugi, Z. Mishal, and T. Tursz. 1985. HLA class-I and class-II antigen expression by human leukemic K562 cells and by Burkitt-K562 hybrids: modulation by differentiation inducers and interferon. Exp Hematol 13: 885–890.

45. Heinzel, A. S., J. E. Grotzke, R. A. Lines, D. A. Lewinsohn, A. L. McNabb, D. N. Streblow, V. M. Braud, H. J. Grieser, J. T. Belisle, and D. M. Lewinsohn. 2002. HLA-E-dependent presentation of Mtb-derived antigen to human CD8+ T cells. The Journal of experimental medicine 196: 1473–1481.

46. core, N. t. 2010. Production Protocols. Emory University.

47. C. Wieczorek, M., E. T. Abualrous, J. Sticht, M. Álvaro-Benito, S. Stolzenberg, F. Noé, and C. Freund. 2017. Major Histocompatibility Complex (MHC) Class I and MHC Class II Proteins: Conformational Plasticity in Antigen Presentation. Front Immunol 8: 292–292.

48. Coppola, C., B. Hopkins, S. Huhn, Z. Du, Z. Huang, and W. J. Kelly. 2020. Investigation of the Impact from IL-2, IL-7, and IL-15 on the Growth and Signaling of Activated CD4(+) T Cells. Int J Mol Sci 21.

49. Bevington, S. L., P. Keane, J. K. Soley, S. Tauch, D. W. Gajdasik, R. Fiancette, V. Matei-Rascu, C. M. Willis, D. R. Withers, and P. N. Cockerill. 2020. IL-2/IL-7-inducible factors pioneer the path to T cell differentiation in advance of lineage-defining factors. The EMBO Journal 39: e105220.

50. Gillis, S., and K. A. Smith. 1977. Long term culture of tumour-specific cytotoxic T cells. Nature 268: 154–156.

51. D. Grant, N. L., P. Maiello, E. Klein, P. L. Lin, H. J. Borish, J. Tomko, L. J. Frye, A. G. White, D. E. Kirschner, J. T. Mattila, and J. L. Flynn. 2022. T cell transcription factor expression evolves over time in granulomas from Mycobacterium tuberculosis-infected cynomolgus macaques. Cell Rep 39: 110826.

52. Ganchua, S. K. C., A. M. Cadena, P. Maiello, H. P. Gideon, A. J. Myers, B. F. Junecko, E. C. Klein, P. L. Lin, J. T. Mattila, and J. L. Flynn. 2018. Lymph nodes are sites of prolonged bacterial persistence during Mycobacterium tuberculosis infection in macaques. PLoS pathogens 14: e1007337.

53. Dimopoulos, N., H. M. Jackson, L. Ebert, P. Guillaume, I. F. Luescher, G. Ritter, and W. Chen. 2009. Combining MHC tetramer and intracellular cytokine staining for CD8+ T cells to reveal antigenic epitopes naturally presented on tumor cells. Journal of Immunological Methods 340: 90–94.

54. Pastore, G., M. Carraro, E. Pettini, E. Nolfi, D. Medaglini, and A. Ciabattini. 2019. Optimized Protocol for the Detection of Multifunctional Epitope-Specific CD4+ T Cells Combining MHC-II Tetramer and Intracellular Cytokine Staining Technologies. Front Immunol 10.

55. Liu, H., M. Rhodes, D. L. Wiest, and D. A. A. Vignali. 2000. On the Dynamics of TCR:CD3 Complex Cell Surface Expression and Downmodulation. Immunity 13: 665–675.

56. Dolton, G., K. Tungatt, A. Lloyd, V. Bianchi, S. M. Theaker, A. Trimby, C. J. Holland, M. Donia, A. J. Godkin, D. K. Cole, P. T. Straten, M. Peakman, I. M. Svane, and A. K. Sewell. 2015. More tricks with tetramers: a practical guide to staining T cells with peptide-MHC multimers. Immunology 146: 11–22.

57. Coppola, M., K. E. van Meijgaarden, K. L. M. C. Franken, S. Commandeur, G. Dolganov, I. Kramnik, G. K. Schoolnik, I. Comas, O. Lund, C. Prins, S. J. F. van den Eeden, G. E. Korsvold, F. Oftung, A. Geluk, and T. H. M. Ottenhoff. 2016. New Genome-Wide Algorithm Identifies Novel In-Vivo Expressed Mycobacterium Tuberculosis Antigens Inducing Human T-Cell Responses with Classical and Unconventional Cytokine Profiles. Scientific reports 6: 37793–37793.

58. O’Connor, S. L., A. J. Blasky, C. J. Pendley, E. A. Becker, R. W. Wiseman, J. A. Karl, A. L. Hughes, and D. H. O’Connor. 2007. Comprehensive characterization of MHC class II haplotypes in Mauritian cynomolgus macaques. Immunogenetics 59: 449–462.

59. Budde, M. L., R. W. Wiseman, J. A. Karl, B. Hanczaruk, B. B. Simen, and D. H. O’Connor. 2010. Characterization of Mauritian Cynomolgus Macaque Major Histocompatibility Complex Class I Haplotypes by High Resolution Pyrosequencing. Immunogenetics 62: 773–780.

60. Brandt, L., Y. A. W. Skeiky, M. R. Alderson, Y. Lobet, W. Dalemans, O. C. Turner, R. J. Basaraba, A. A. Izzo, T. M. Lasco, P. L. Chapman, S. G. Reed, and I. M. Orme. 2004. The protective effect of the Mycobacterium bovis BCG vaccine is increased by coadministration with the Mycobacterium tuberculosis 72-kilodalton fusion polyprotein Mtb72F in M. tuberculosis-infected guinea pigs. Infection and immunity 72: 6622–6632.

61. B. Tsenova, L., R. Harbacheuski, A. L. Moreira, E. Ellison, W. Dalemans, M. R. Alderson, B. Mathema, S. G. Reed, Y. A. W. Skeiky, and G. Kaplan. 2006. Evaluation of the Mtb72F polyprotein vaccine in a rabbit model of tuberculous meningitis. Infection and immunity 74: 2392–2401.

62. C. Reed, S. G., R. N. Coler, W. Dalemans, E. V. Tan, E. C. DeLa Cruz, R. J. Basaraba, I. M. Orme, Y. A. W. Skeiky, M. R. Alderson, K. D. Cowgill, J.-P. Prieels, R. M. Abalos, M.- C. Dubois, J. Cohen, P. Mettens, and Y. Lobet. 2009. Defined tuberculosis vaccine, Mtb72F/AS02A, evidence of protection in cynomolgus monkeys. Proceedings of the National Academy of Sciences 106: 2301–2306.

63. Von Eschen, K., R. Morrison, M. Braun, O. Ofori-Anyinam, E. De Kock, P. Pavithran, M. Koutsoukos, P. Moris, D. Cain, M. C. Dubois, J. Cohen, and W. R. Ballou. 2009. The candidate tuberculosis vaccine Mtb72F/AS02A: Tolerability and immunogenicity in humans. Hum Vaccin 5: 475–482.

64. Leroux-Roels, I., G. Leroux-Roels, O. Ofori-Anyinam, P. Moris, E. De Kock, F. Clement, M. C. Dubois, M. Koutsoukos, M. A. Demoitié, J. Cohen, and W. R. Ballou. 2010. Evaluation of the safety and immunogenicity of two antigen concentrations of the Mtb72F/AS02(A) candidate tuberculosis vaccine in purified protein derivative-negative adults. Clin Vaccine Immunol 17: 1763–1771.

65. Griffiths, K. L., A. A. Pathan, A. M. Minassian, C. R. Sander, N. E. R. Beveridge, A. V. S. Hill, H. A. Fletcher, and H. McShane. 2011. Th1/Th17 Cell Induction and Corresponding Reduction in ATP Consumption following Vaccination with the Novel Mycobacterium tuberculosis Vaccine MVA85A. PLOS ONE 6: e23463.

66. A. Scriba, T. J., B. Kalsdorf, D.-A. Abrahams, F. Isaacs, J. Hofmeister, G. Black, H. Y. Hassan, R. J. Wilkinson, G. Walzl, S. J. Gelderbloem, H. Mahomed, G. D. Hussey, and W. A. Hanekom. 2008. Distinct, Specific IL-17- and IL-22-Producing CD4&lt;sup&gt;+&lt;/sup&gt; T Cell Subsets Contribute to the Human Anti-Mycobacterial Immune Response. The Journal of Immunology 180: 1962.

67. Kallies, A., and K. L. Good-Jacobson. 2017. Transcription Factor T-bet Orchestrates Lineage Development and Function in the Immune System. Trends in Immunology 38: 287–297.

68. Dijkman, K., N. Aguilo, C. Boot, S. O. Hofman, C. C. Sombroek, R. A. W. Vervenne, C. H. M. Kocken, D. Marinova, J. Thole, E. Rodríguez, M. P. M. Vierboom, K. G. Haanstra, E. Puentes, C. Martin, and F. A. W. Verreck. 2021. Pulmonary MTBVAC vaccination induces immune signatures previously correlated with prevention of tuberculosis infection. Cell Rep Med 2: 100187.

69. J. Gideon, H. P., T. K. Hughes, C. N. Tzouanas, M. H. Wadsworth, 2nd, A. A. Tu, T. M. Gierahn, J. M. Peters, F. F. Hopkins, J. R. Wei, C. Kummerlowe, N. L. Grant, K. Nargan, J. Y. Phuah, H. J. Borish, P. Maiello, A. G. White, C. G. Winchell, S. K. Nyquist, S. K. C. Ganchua, A. Myers, K. V. Patel, C. L. Ameel, C. T. Cochran, S. Ibrahim, J. A. Tomko, L. J. Frye, J. M. Rosenberg, A. Shih, M. Chao, E. Klein, C. A. Scanga, J. Ordovas-Montanes, B. Berger, J. T. Mattila, R. Madansein, J. C. Love, P. L. Lin, A. Leslie, S. M. Behar, B. Bryson, J. L. Flynn, S. M. Fortune, and A. K. Shalek. 2022. Multimodal profiling of lung granulomas in macaques reveals cellular correlates of tuberculosis control. Immunity 55: 827–846 e810.

70. Kauffman, K. D., S. Sakai, N. E. Lora, S. Namasivayam, P. J. Baker, O. Kamenyeva, T. W. Foreman, C. E. Nelson, D. Oliveira-de-Souza, C. L. Vinhaes, Z. Yaniv, C. S. Lindestam Arleham, A. Sette, G. J. Freeman, R. Moore, N. D. T. I. Program, A. Sher, K. D. Mayer-Barber, B. B. Andrade, J. Kabat, L. E. Via, and D. L. Barber. 2021. PD-1 blockade exacerbates &lt;em&gt;Mycobacterium tuberculosis&lt;/em&gt; infection in rhesus macaques. Science Immunology 6: eabf3861.

71. Basdeo, S. A., D. Cluxton, J. Sulaimani, B. Moran, M. Canavan, C. Orr, D. J. Veale, U. Fearon, and J. M. Fletcher. 2017. Ex-Th17 (Nonclassical Th1) Cells Are Functionally Distinct from Classical Th1 and Th17 Cells and Are Not Constrained by Regulatory T Cells. J Immunol 198: 2249–2259.

72. Cibrián, D., and F. Sánchez-Madrid. 2017. CD69: from activation marker to metabolic gatekeeper. Eur J Immunol 47: 946–953.

73. González-Amaro, R., J. R. Cortés, F. Sánchez-Madrid, and P. Martín. 2013. Is CD69 an effective brake to control inflammatory diseases? Trends Mol Med 19: 625–632.

